# Rare CNVs and phenome-wide profiling: a tale of brain-structural divergence and phenotypical convergence

**DOI:** 10.1101/2022.04.23.489093

**Authors:** J. Kopal, K. Kumar, K. Saltoun, C. Modenato, C. A. Moreau, S. Martin-Brevet, G. Huguet, M. Jean-Louis, C.O. Martin, Z. Saci, N. Younis, P. Tamer, E. Douard, A. M. Maillard, B. Rodriguez-Herreros, A. Pain, S. Richetin, 16p11.2 European Consortium, Simons Searchlight Consortium, L. Kushan, A. I. Silva, M. B. M. van den Bree, D. E. J. Linden, M. J. Owen, J. Hall, S. Lippé, B. Draganski, I. E. Sønderby, O. A. Andreassen, D. C. Glahn, P. M. Thompson, C. E. Bearden, S. Jacquemont, D. Bzdok

## Abstract

Copy number variations (CNVs) are rare genomic deletions and duplications that can exert profound effects on brain and behavior. Previous reports of pleiotropy in CNVs imply that they converge on shared mechanisms at some level of pathway cascades, from genes to large-scale neural circuits to the phenome. However, studies to date have primarily examined single CNV loci in small clinical cohorts. It remains unknown how distinct CNVs escalate the risk for the same developmental and psychiatric disorders. Here, we quantitatively dissect the impact on brain organization and behavioral differentiation across eight key CNVs. In 534 clinical CNV carriers from multiple sites, we explored CNV-specific brain morphology patterns. We extensively annotated these CNV-associated patterns with deep phenotyping assays through the UK Biobank resource. Although the eight CNVs cause disparate brain changes, they are tied to similar phenotypic profiles across ∼1000 lifestyle indicators. Our population-level investigation established brain structural divergences and phenotypical convergences of CNVs, with direct relevance to major brain disorders.

## Introduction

A chief goal of modern neuroscience is understanding how genetic variation impacts brain organization and inter-individual differences in behavior. Advances in genomic microarray technology streamlined the detection of copy number variations (CNVs) – deletions or duplications of chromosomal segments of >1000 base pairs (Conrad et al., 2010; Freeman et al., 2006). This class of genetic mutations opens a unique window into the investigation of how neurogenetic determinants shape human behavior, cognition, and development (Auwerx et al., 2022; Jacquemont et al., 2021). Pathogenic CNVs, consistently reoccurring when considering many individuals, provide opportunities to study groups of individuals who carry the same deletion or duplication of a well-defined set of genes (Rutkowski et al., 2017). Moreover, CNVs affecting multiple genes have larger effects on phenotype than the low effect-size single-nucleotide polymorphisms identified by genome-wide association studies (Lauer and Gresham, 2019). Concretely, CNVs overall have been shown to detrimentally affect cognition and raise the risk for psychiatric conditions (Huguet et al., 2021; Jacquemont et al., 2021). Nevertheless, the nature and specificity of CNV effects on phenotypic traits have become an arena of ongoing investigation (Jacquemont et al., 2021; Moberg et al., 2018; Silva et al., 2022).

CNVs can serve as a tool to scrutinize genetic mechanisms underlying vulnerability to brain disorders in general and neuropsychiatric disorders in particular, irrespective of clinical symptomatology. The vast majority of large recurrent CNVs have been linked to more than one clinical diagnosis including intellectual disability, autism spectrum disorders, and schizophrenia (Chawner et al., 2019; Marshall et al., 2017; Sanders et al., 2019). These findings make a case that circumscribed genetic changes are rarely exclusively associated with a single clinical diagnosis (Moreno-De-Luca and Martin, 2021). Further, CNVs have demonstrable consequences even in seemingly unaffected middle and old age carriers, who show no overt signs of early-onset neuropsychiatric disorders. Specifically, recent evidence points to a wider spectrum of impacts from CNV status, ranging from physical traits to diabetes to hypertension to obesity to renal dysfunction (Auwerx et al., 2022; Crawford et al., 2019; Owen et al., 2018) as well as psychopathology (Adams et al., 2022). Understudied body-wide CNV effects may contribute to the links of schizophrenia-associated CNVs with diminished academic qualifications, occupation, or household income (Kendall et al., 2019). In summary, this class of genetic variants can be associated with an overlapping set of behavioral and clinical phenotypes, despite affecting distant and disparate parts of the genome (Doelken et al., 2013; Viñas-Jornet et al., 2018). Nevertheless, why so many CNVs increase the risk for the same developmental and psychiatric disorders remain unexplained.

Despite many advances in genomic profiling, investigations into the corresponding brain signatures have only been performed for a few CNVs and mostly focused on a single variant at a time (Moreau et al., 2020, 2021b). These parallel approaches to catalog CNVs highlighted a wide spectrum of robust effects on brain structure (Modenato et al., 2021b; Sønderby et al., 2021). Nevertheless, even though distinct rare CNVs are associated with a range of brain alterations, they have been argued to lead to a degree of similarity in associated behavioral phenotypes (Raznahan et al., 2022; Silva et al., 2022). However, it is unknown at which level of observation, from genes to large-scale brain networks to the phenome, the mechanisms of this disparate group of risk variants converge. Since deleterious CNVs are rare, such as 1 in 3,000 for 22q11.2 deletion (Zinkstok et al., 2019), previous investigations suffered from small samples of subjects and a lack of phenotypic depth. Therefore, previous studies were chronically underpowered to paint a complete picture of CNVs in neuroscience, particularly how associated brain-imaging signatures may relate to clinical phenotypes. There is a need for a systematic investigation of intermediate brain measures and quantitative phenotypic measures that will ultimately help determine at what level convergence occurs from genes to phenotypes. Consequently, the recent advent of population cohorts with rich phenotypic assessment batteries represents an untapped opportunity to conjointly examine a set of CNVs and characterize them at an unprecedented scale.

Population-based cohorts are ideally suited to tease apart the commonalities in phenotypic outcomes across CNV alterations. In the present study, we interrogated the largest existing biomedical data resource, the UK Biobank, which allowed a head-to-head comparison of an envelope of CNVs. Specifically, we leveraged tools from machine learning, such as linear discriminant analysis (LDA), to isolate CNV-specific brain morphology signatures from a multisite clinical cohort. These individuals carried one of eight rare recurrent CNVs previously associated with developmental brain disorders that have been commonly studied in isolated efforts. Subsequently, we tailored an analytic strategy to carry over the specific CNV brain signatures from the clinically ascertained cohort to the large-scale UK Biobank cohort. We directly linked a rich portfolio of phenotypes to the presence of the eight CNV brain signatures in ∼40,000 UK Biobank participants. Concretely, capitalizing on the largest brain-imaging CNV dataset to date and building on the breadth of phenotypic annotation available in the UK Biobank, we performed separate phenome-wide association studies (PheWAS) for the eight CNV-brain-imaging signatures across 977 phenotypes spanning eleven categories. In this way, we provide a population-level characterization of what unites and divides the eight recurrent CNVs by detailing convergences and divergences from genomic variants to brain morphology to phenome. In an attempt to establish cornerstone evidence for the community, such a study can illuminate fundamental links between genetic variation and brain organization, with their consequences for behavioral manifestations.

## Results

### Dissecting different CNV effects on whole-brain morphology

We systematically analyzed volumetric measures derived from brain-imaging scans in the clinical cohort comprising 846 total subjects: 534 carried one of eight recurrent CNVs (deletion and duplications of 1q21.1 distal, 15p11.2, 16p11.2 proximal, or 22q11.2), while 312 controls did not carry a CNV (Tab. 1). We parsed volume measures from these structural brain scans using a 400-region anatomical definition (Schaefer-Yeo reference atlas; see Online methods). To account for variation outside of our current primary scientific interest, each brain region volume was adjusted for total grey matter, age, age^2^, sex, and acquisition site for all downstream analysis steps.

In a first step, separately for each of the eight CNVs, we quantified their effects on brain region volume measures. Specifically, after normalizing (z-scoring) brain volumes across subjects, we examined the extent of volumetric divergence between carriers of each single CNV and controls by computing Cohen’s *d* (giving an effect size for the group difference) for each individual brain region (Fig. 1a). In doing so, for each examined CNV, we obtained a brain map of Cohen’s *d* effect sizes that summarize magnitudes of CNV-induced structural abnormalities across the brain’s grey matter. We noted widespread smaller volumes in the majority of the examined atlas regions for the 1q21.1 distal deletion, the 15q11.2 duplication, the 16p11.2 proximal duplication, and the 22q11.2 deletion. Conversely, a preferential increase in most regional volumes became apparent for the 1q21.1 distal duplication, 15q11.2 deletion, 16p11.2 proximal deletion, and the 22q11.2 duplication. As such, each target CNV locus was characterized by an overall constellation of grey matter changes – a brain-wide CNV map of how particular CNV carriership results in systematic brain deviations from controls.

**Figure 1.**
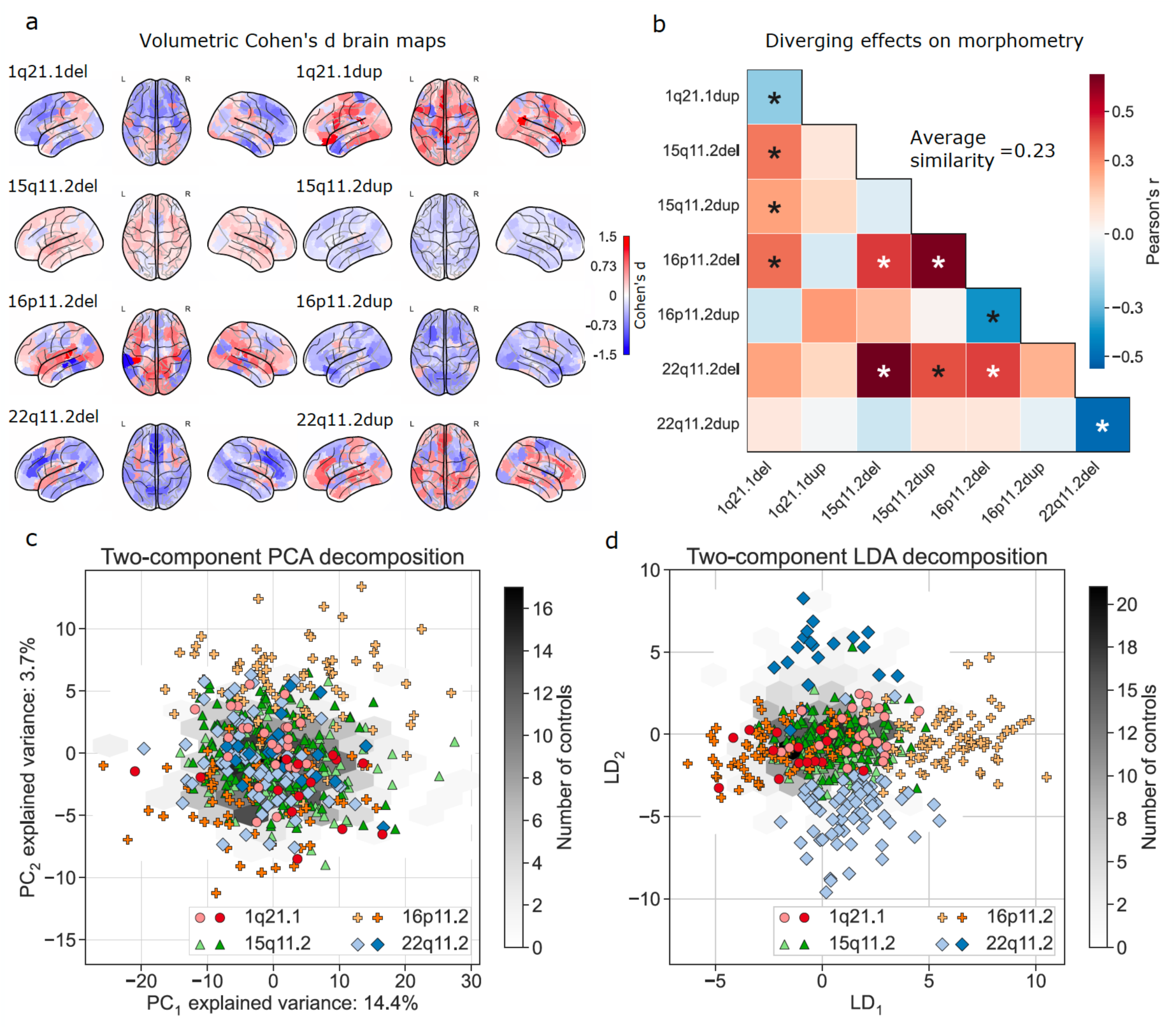
Eight CNVs lead to largely distinct spatial patterns of abnormalities in brain morphology. In 534 subjects carrying one of eight rare CNVs, we have computed Cohen’s d between the carriers and 312 controls, separately for each of the 400 brain regions (Schaefer-Yeo reference atlas). Grey matter region volumes were adjusted for total grey matter, age, age^2^, sex, and acquisition site. a) Cohen’s *d* brain map quantifies the magnitude of structural change for each CNV. Our analysis reveals increased (red) and decreased (blue) brain volumes depending on the variation type. b) Examining associations between the Cohen’s *d* brain maps rendered on brain surface from each pair of CNVs. The wide range and low magnitude of Pearson’s correlations show that CNVs have distinct effects on brain volumes (more red=more similar, more blue=more dissimilar). Average similarity stands for the mean absolute Pearson’s correlations across all CNVs. 22q11.2 and 16p11.2 proximal deletions and duplications show strong mirroring (opposing) effects. Asterisk denotes FDR-corrected spin permutation p-values. c) Projections of z-scored (across subjects) brain volumes onto two dominant dimensions of variation using principal component analysis (PCA). Although the first two dominant PCA components explain 18 % of the variance, they are not related to differences between CNVs. The light and dark symbols represent deletions and duplication, respectively. The grey hexagonal bin plot represents the frequency of controls. Controls were not used to calculate the PCA and were projected post hoc. d) Projections of z-scored (across subjects) brain volumes to two dimensions using linear discriminant analysis (LDA). The first LDA dimension (LD_1_) mainly captures differences between 16p11.2 proximal deletion and duplication, while the second LDA dimension (LD_2_) mainly captures differences between 22q11.2 deletions and duplications. Symbols and hexagonal binning plots were constructed in the same way as for the PCA approach. CNVs lead to distinct changes often represented by a predominant increase or decrease in the grey matter cortex that could effectively be described using low dimensional representations derived by LDA models (cf. methods).

Taken together, this cursory analysis indicated that spatial distributions of mutation-induced changes in brain morphology differed considerably across CNVs (Fig. 1b). To delineate the similarity among effect-size brain maps, we computed Pearson’s correlation between all 400 regional Cohen’s *d* values corresponding to each pair of CNVs. Statistical significance was assessed using a spin-permutation test across the whole brain surface. We found a large disparity between Cohen’s d maps evidenced by the wide spectrum of Pearson’s correlations ranging from -0.51 to 0.63. Concretely, we noted certain similarities, such as for deletions of 22q11.2 and 15q11.2 (r = 0.66, p_FDR-adj_ = 0.03). Further, we observed a strong mirroring effect with significant anti-correlations between deletions and duplication of reciprocal loci. Mirroring effects were strongest for 22q11.2 (r = -0.51, p_FDR-adj_ = 0.03), followed by 16p11.2 proximal (r = -0.39, p_FDR-adj_ = 0.03). The average volumetric similarity measured by the average absolute Pearson’s correlations was r = 0.23. Together, we quantified the diverging effects of CNV loci on the human cortex.

### Visualizing CNV differences in low-dimensional summary signatures

A drawback of the approach based on Cohen’s *d* lies in its univariate character, which considers each region separately, ignoring the respective remaining atlas regions. Hence, next, we used dimensionality reduction techniques to obtain holistic summaries of the CNV carriers’ morphological profiles. We set out from the possibility that CNVs cause coordinated volume changes distributed across the entire brain. Therefore, we expected an intrinsically brain-spanning pattern could be extracted that faithfully captures the induced morphological differences. Principal component analysis (PCA) is the most commonly used multivariate tool that is demonstrably most effective at representing linear latent factors. Concretely, PCA can be interpreted as computing a new coordinate system such that the axes are oriented in the directions of the largest variation across the 400 region volume measures. We thus used PCA to project all CNV carriers’ regional volumes (normalized based on distribution from controls) onto the two dominant directions of coherent whole-cortex variation (Fig. 1c). In the ensuing two-dimensional subject embedding, CNV carriers were scattered randomly without an apparent systematic relationship with each other. In other words, the results suggested that CNVs were not the primary source of the interindividual variation in whole-cortex morphology in our cohort. Hence, a method without access to CNV-carriership status, such as PCA, could not provide a satisfying overall description of what drives structural brain deviations induced by specific CNV loci.

Therefore, we turned to linear discriminant analysis (LDA) as a pattern classification algorithm that is naturally capable of recovering a low-dimensional representation explicitly aimed at maximizing the separation between the eight CNVs based on the individuals’ brain morphometry measures. We then re-expressed the brain-wide regional volumes as the two primary dimensions of structural variation under the LDA model (Fig. 1d). In particular, the leading dimension of the LDA-derived subject embedding captured the differences between 16p11.2 proximal deletion and duplications. The second most explanatory dimension of the LDA-derived embedding mainly captured the differences between 22q11.2 deletion and duplications. This distribution of a single CNV locus along a single dimension points again at similar structural effects with opposite directions. In summary, LDA formed a new low-dimensional space in which the brain morphology of CNV carriers could be effectively identified, quantified, and, subsequently, examined in further detail.

### Deriving intermediate phenotypes that track subject-specific expressions of CNV signatures

To supplement the multi-CNV classification model which explored differences between CNV loci (described above), our next analysis step was to bring to light differences between carriers of a given CNV and controls. Therefore, we constructed eight LDA models dedicated to the eight CNVs. Notably, there was a considerable imbalance between the number of controls and CNV carriers (from 2-fold for 15q11.2 duplication to 22-fold for 1q21.1 distal duplication). Moreover, the number of model parameters to be estimated (at least 400 parameters associated with the 400 atlas regions) was larger than the number of subjects. To remedy the challenges of this data scenario, our analysis pipeline combined bagging, bootstrapping, and regularization to prevent overfitting the model hyperparameters (see details in Online methods). We evaluated the model performance indexed by out-of-sample prediction in MRI brain scans unseen by the model using Matthews correlation coefficient. All CNVs were successfully classified with a consistent above-chance accuracy (Fig. 2a). Chance level accuracy was defined as the performance of an empirical null model obtained by label shuffling. Furthermore, the classification performance served as one possible proxy to quantify the divergence of CNV carriers from the population. In other words, 22q11.2 and 16p11.2 proximal deletion carriers emerged to yield a particularly distinct brain pattern, while 15q11.2 duplications were relatively more similar to the general population (Fig. 2a).

**Figure 2.**
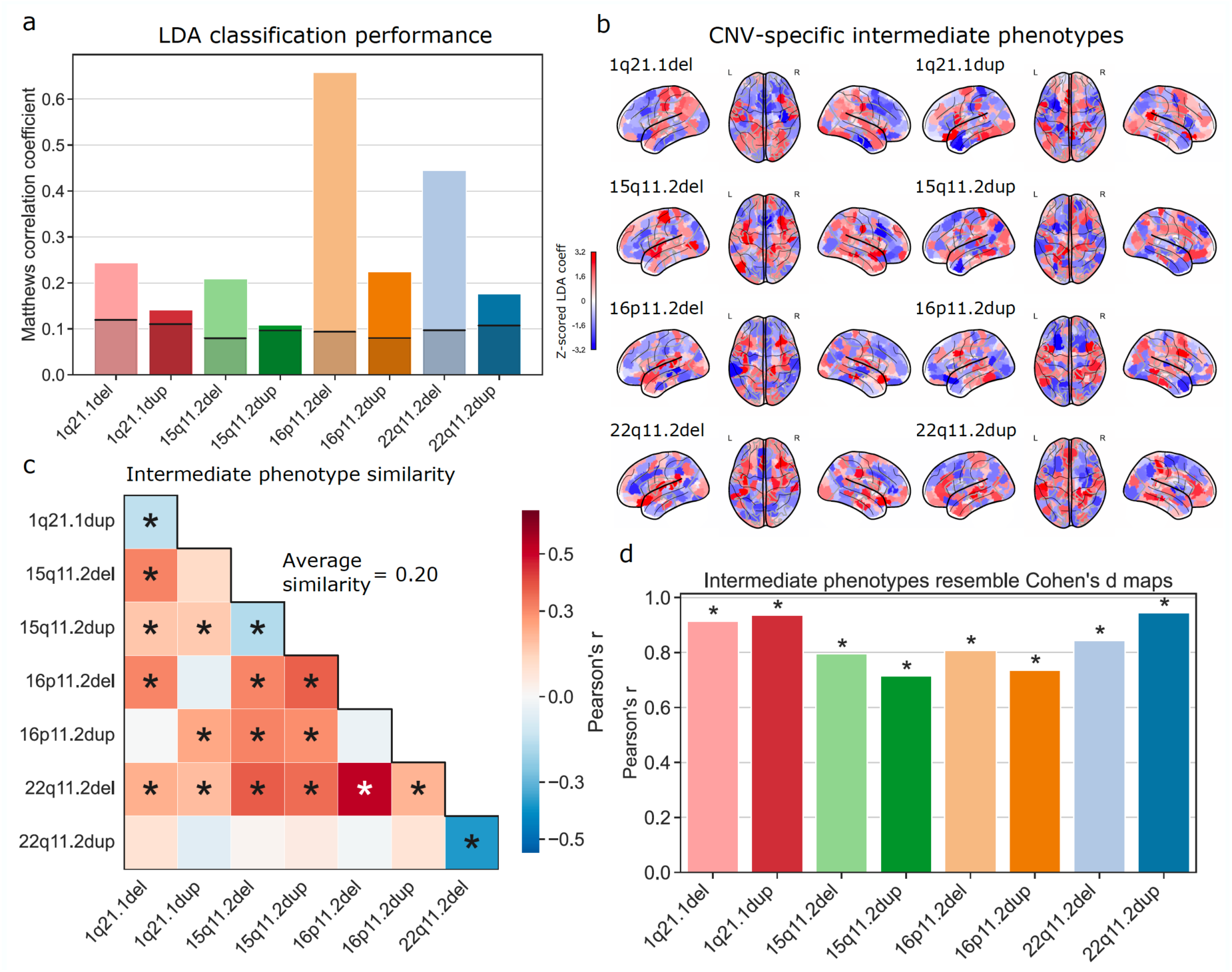
Pattern-learning models extract distinct intermediate brain phenotypes from CNV status. We estimated eight LDA models to classify between controls and each of the eight different CNVs. Regularization, bootstrapping, and bagging ensured that the learning algorithms did not overfit (cf. Online methods). a) Classification performance of eight distinct LDA models when telling apart controls and CNV carriers, given as Matthews correlation coefficient. All eight CNVs are successfully classified based on brain structure, at above-chance accuracy as their performance exceeds that of an empirical null model (95% confidence intervals shown, obtained by label shuffling). b) Shows the prediction rule derived for each of the eight CNV-specific LDA models projected on the brain (red/blue = positive/negative weight). The prediction rule is a CNV-specific brain signature and can be treated as an intermediate phenotype. c) Similarity between CNV-specific intermediate phenotypes. The wide range and low magnitudes of ensuing Pearson’s correlations reflect the disparity in the captured intermediate phenotypes. Average similarity represents the mean absolute correlation across all CNVs. Asterisk denotes FDR-corrected spin permutation p-values. d) Relationship between Cohen’s *d* brain maps and intermediate phenotypes. All eight intermediate phenotypes appear to largely follow the respective Cohen’s *d* brain maps. LDA models identified and quantified CNV-specific intermediate phenotypes that effectively captured distinct morphometric differences between CNV carriers and the general population.

After extracting predictive principles of structural brain deviations by means of LDA, each formed model included a collection of 400 coefficients associated with the atlas regions (Fig. 2b). These coefficients encapsulated a multivariate prediction rule which maximized the difference between controls and CNV carriers. In other words, each CNV’s LDA model encapsulated an intermediate phenotype – a brain-wide volumetric signature that characterizes each CNV. To quantify the similarity between the derived intermediate phenotype representations, we compared them using Pearson’s correlation coefficient. Again, we observed certain similarities across the eight CNVs, as well as mirroring effects between reciprocal CNVs (Fig. 2c). However, the wide range and low strength (average similarity r = 0.2) of obtained CNV-CNV similarities indicated that LDA models reflected the sizable diverging effects of CNVs on brain morphometry. In concordance, the identified intermediate phenotypes bore a degree of similarity to the Cohen’s d brain maps (Fig. 2d). The strong positive Pearson’s correlation between the intermediate phenotypes and Cohen’s *d* brain maps was significant for all CNVs. As such, our approach allowed us to extract CNV-specific intermediate phenotypes that effectively captured volumetric differences between CNV carriers and controls.

We further inspected the 400 region coefficients of each LDA model that captured the influence of each specific CNV on each brain region. By carrying out a one-sample bootstrap hypothesis test independently for each CNV, we assessed which region-specific model coefficients are robustly different from zero and thus robustly affected by CNVs. During the learning process, we reiterated each CNV-specific LDA model 100 times; every time with a different set of subjects based on resampling the 846 subjects with repetition. Statistically relevant coefficients were robustly different from zero if their two-sided confidence interval - according to the 2.5/97.5% intervals of the bootstrap-derived distribution - did not include zero. Different CNVs affected (displayed statistically relevant coefficients) different layers that correspond to the seven large-scale brain networks populating the cortex, as defined by our atlas (Fig. 3a). For example, while 16p11.2 proximal duplication primarily affects 20% of all regions in the limbic network, 22q11.2 deletion affects 20% of regions in the salience ventral attentional network as well as more than 10% of regions in the limbic, dorsal attentional, and default-mode networks. Across all examined CNVs and target brain networks, the 16p11.2 proximal deletion affected the largest number of brain regions, while 15q11.2 duplication affected the lowest number of regions. Higher-order network circuits showed, on average, the highest number of relevant coefficients. Concretely, the limbic network had the highest relative number of affected regions, followed by the salience and default-mode networks (Fig. 3b). Altogether, the coefficients spanning several large-scale networks formed the backbone of each CNV’s intermediate phenotype.

**Figure 3.**
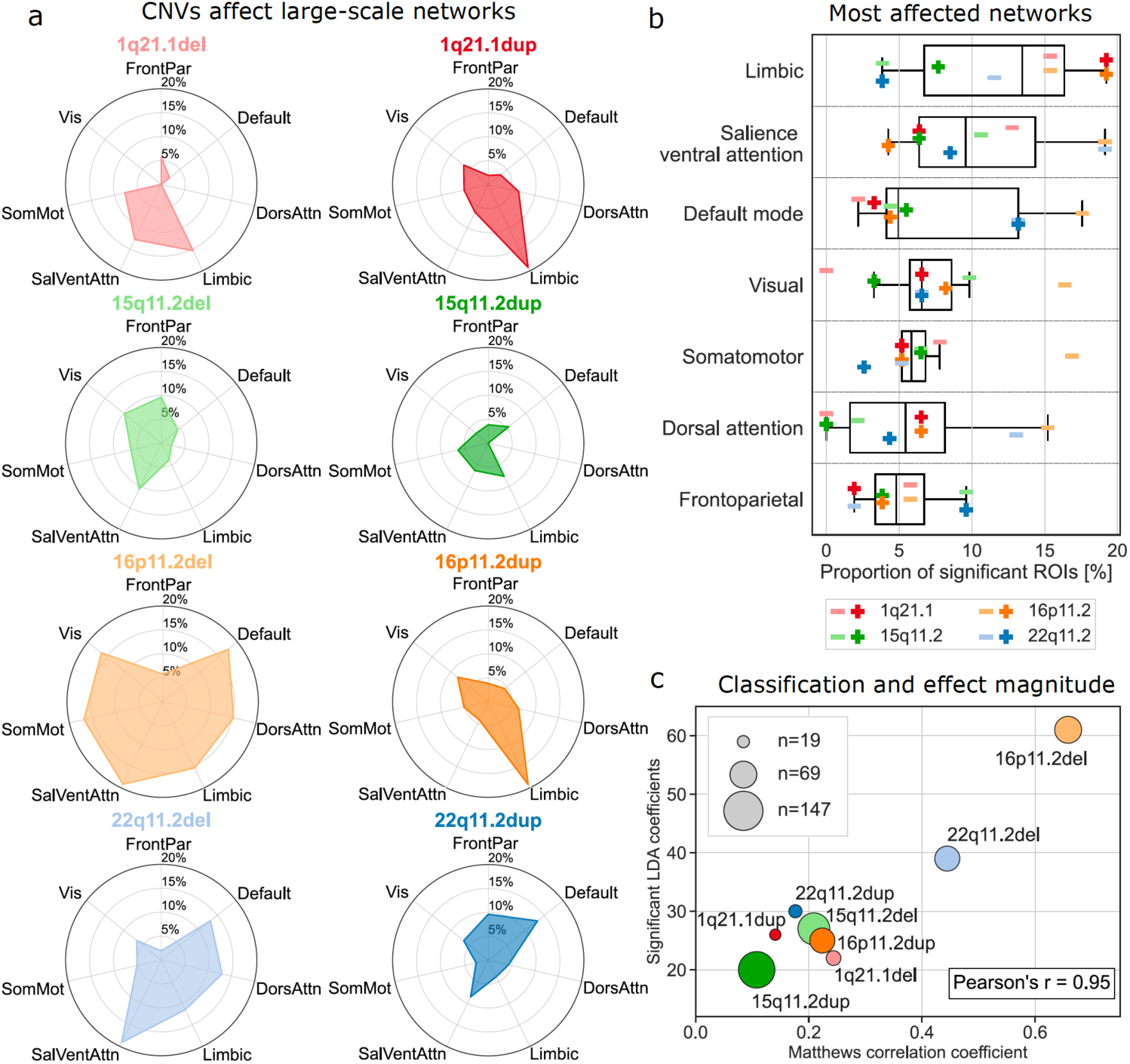
Intermediate brain phenotypes track structural changes with distinct impacts in large-scale networks. We identified which aspects of the LDA-derived prediction rule robustly contributed to classification success by calculating 100 bootstrapped LDA models for each CNV while sampling CNV carriers randomly. a) Percentage of statistically relevant LDA coefficients in a CNV carrier group among all the regions that belong to each brain network (one-sample bootstrap hypothesis test for non-zero mean with 10,000 replicates). For example, 16p11.2 proximal deletion strongly affects most large-scale networks except the frontoparietal network. Altogether, the estimated LDA coefficients represent the backbone of each intermediate phenotype. Large-scale networks correspond to seven SchaeferYeo networks; Vis: Visual, FrontPar: Frontoparietal, SomMot: Somatomotor, DorsAttn: Dorsal attention, SalVenAttn: Salience ventral attention, Limbic, Frontoparietal, Default: Default mode. b) LDA coefficients grouped by the large-scale networks. The highest relative number of affected regions is in the limbic network. Conversely, regions in the frontoparietal network are targeted less frequently. c) Relationship between CNV effects and LDA performance. There is a significant positive correlation between the number of significant LDA coefficients and classifier performance, unlike for the sample size of the cohort (marker size). According to the eight specific LDA models, CNVs affected predominantly high-level networks such as the limbic, salience, and default-mode networks.

To further explore characteristic relationships between the eight CNVs, we probed for a linear relationship of the number of salient LDA coefficients with LDA classifier performance and average brain-wide Cohen’s *d* (see above). We found significant positive Pearson’s correlation with classifier performance (r = 0.95, p = 0.0003) (Fig. 3c) while the linear relationship with mean absolute effect size was not significant (r = 0.47, p = 0.24). Furthermore, when we included sample size in the testing scheme, we found only a negative linear association with average Cohen’s *d* (r = -0.80, p = 0.02), calling for careful interpretation of effect sizes, owing to the estimation of population mean in small samples. In sum, our collective findings highlighted how LDA models reflect CNV-specific changes in large-scale brain networks to form distinctive intermediate phenotypes.

### Lifting over the intermediate phenotypes from the clinical cohort to the UK Biobank

We built eight separate LDA models that encapsulated CNV-specific intermediate phenotypes. By doing so, we could quantify the presence of each intermediate CNV phenotype for each subject. Hence, as an illustrative example, we compared the expression level of 16p11.2 proximal duplication intermediate phenotype between the carriers of that CNV and controls. Based on a two-sample bootstrap hypothesis test for difference of means with 10,000 bootstrap iterations (Online methods), the ensuing means of the intermediate phenotype expressions differed robustly between CNV carriers in the clinical sample and controls (p-value < 10^−4^) (Fig. 4a). Critically, the predictive rule from each CNV-specific LDA model derived from our clinical population could be used to quantify expressions of that intermediate phenotype in a phenotypically richer population data repository.

**Figure 4.**
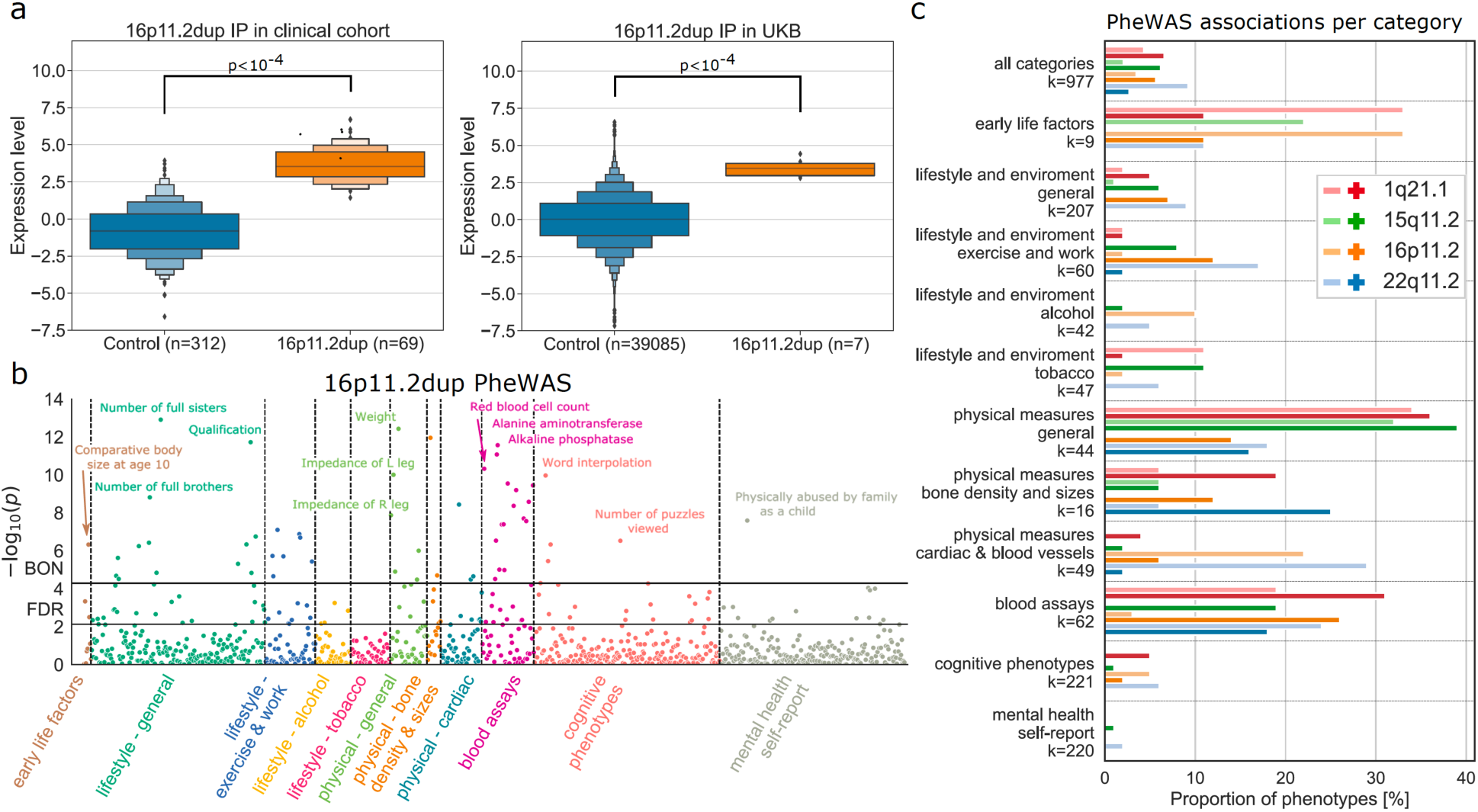
Schematic: Using intermediate CNV phenotypes as a basis for phenome-wide association analysis. We performed a phenome-wide association study (PheWAS) by computing Pearson’s correlation between the expression of each of the eight intermediate CNV phenotypes and 977 phenotypes spanning 11 categories in 39,085 UK Biobank subjects. a) Letter-value (boxen) plot for the expression of 16p11.2 proximal duplication intermediate phenotype is shown for the sake of illustration. These expression scores are computed by quantifying the presence of derived 16p11.2 proximal duplication intermediate phenotype in both the clinical cohort (left) and the UK Biobank (right). Based on a two-sample bootstrap hypothesis test for difference of means with 10,000 bootstrap replicates, the 16p11.2 proximal duplication carriers significantly differed in the expression level from controls both in the clinical cohort (p <10^−4^) and UK Biobank dataset (p <10^−4^) b) PheWAS study using the CNV-specific intermediate phenotype derived by LDA. We calculated the Pearson’s correlation between the expression of 16p11.2 proximal duplication intermediate phenotype and each of 977 phenotypes. After the Bonferroni correction for multiple comparisons (BON), there were 55 significant associations, such as education score, haemoglobin concentration, or physically abused by family as a child. There were 145 significant associations exceeding false discovery rate correction (FDR) c) The relative number of significant correlations summarized for each of the eleven categories for each CNV in the UK Biobank. Most CNVs are strongly associated with multiple categories and their respective phenotypes. For example, up to 35% of phenotypes in general physical measures show a significant correlation with four CNVs brain signatures. The light and dark symbols represent deletions and duplication, respectively. As an insight from the performed phenome-wide association analysis, CNV brain signatures are linked with multiple phenotypes across most categories but mainly in the general physical measures, blood assays, and early life factors categories.

To carry over the intermediate phenotypes from the clinical cohort to the UK Biobank, we quantified the expression of each intermediate CNV phenotype for all 39,085 UK Biobank participants. Concretely, we first extracted brain volume measures from the 400 atlas regions, adjusting for several confound variables (see Online methods). We then calculated the subject-specific expression for all intermediate CNV phenotypes in the UK Biobank. The generalizability of the derived intermediate phenotypes was indicated by the difference between the intermediate phenotype expression level of non-carriers and CNV carriers in the UK Biobank (for 16p11.2 proximal duplication p-value < 10^−4^ using an identical test to that above, Fig. 4a). Notably, we obtained similar results for all other seven intermediate phenotypes (Supp. Fig. 1). Carrying over CNV-associated MRI profiles computed in the clinical cohort to the UK Biobank was a critical step that allowed us to identify phenotype correlates of the CNV-associated MRI profiles in a population >500 times larger than our median CNV cohort.

### Charting phenome-wide associations of CNV phenotype expression in the UK Biobank

The UK Biobank is the largest existing uniform brain-imaging dataset in terms of subject sample size and the breadth of available phenotypic annotations. Building on the derived subject-specific expressions of the eight intermediate CNV phenotypes, we performed a phenome-wide association study (PheWAS) by calculating Pearson’s correlation between the intermediate phenotype expression and each of the 977 unique phenotypes provided by the UK Biobank resource (Fig. 4b), spanning eleven different categories (Supp. Fig. 2a). In our recurring example of the 16p11.2 proximal duplication intermediate phenotype, 55 associations surpassed Bonferroni correction for multiple testing (including comparative body size at age 10, education score, haemoglobin concentration, or physically abused by family as a child), while 145 associations surpassed FDR correction. In other words, individuals with greater similarity to the 16p11.2 proximal duplication MRI profiles showed a stronger association with levels of education or blood assays biomarkers.

To gain additional insight, we summarized the phenotypic association profiles by domain. To this end, we calculated the relative number of association hits for each of the eleven phenotypic domains (using the more stringent Bonferroni correction) as a ratio between the number of significant associations and the number of phenotypes in each category. The highest relative number of associations were in categories detailing physical measures, blood assays, and early life factors categories (Fig. 4c). The collective results showed that CNVs are associated with numerous rich and diverse phenotypes across all eleven categories.

Among all examined CNVs, the 22q11.2 deletion intermediate phenotype displayed the highest number of phenome-wide hits, with 90 robust associations after Bonferroni’s correction for multiple comparisons (Fig. 4c; for further details, see Supp. Fig. 3-10). Furthermore, we examined the similarity of phenotypic profiles across CNVs, analogous to comparing volumetric signatures (cf. above). To this end, we calculated a correlation between the association strengths (Pearson’s correlations) from each PheWAS analysis (Fig. 5a). The definitive collection of brain signature-phenotype links reflected the linear association strength between CNV phenotypical profiles across 977 indicators (Fig. 5b). We found a strong resemblance (average similarity r = 0.62) between the eight phenotypical profiles with positive as well as negative correlations (Pearson’s correlations from r = -0.84 to 0.82). In other words, although CNVs displayed different brain morphometric effects, their volumetric signatures appeared to be linked with similar phenotypes across a rich portfolio of ∼1000 curated lifestyle indicators.

**Figure 5.**
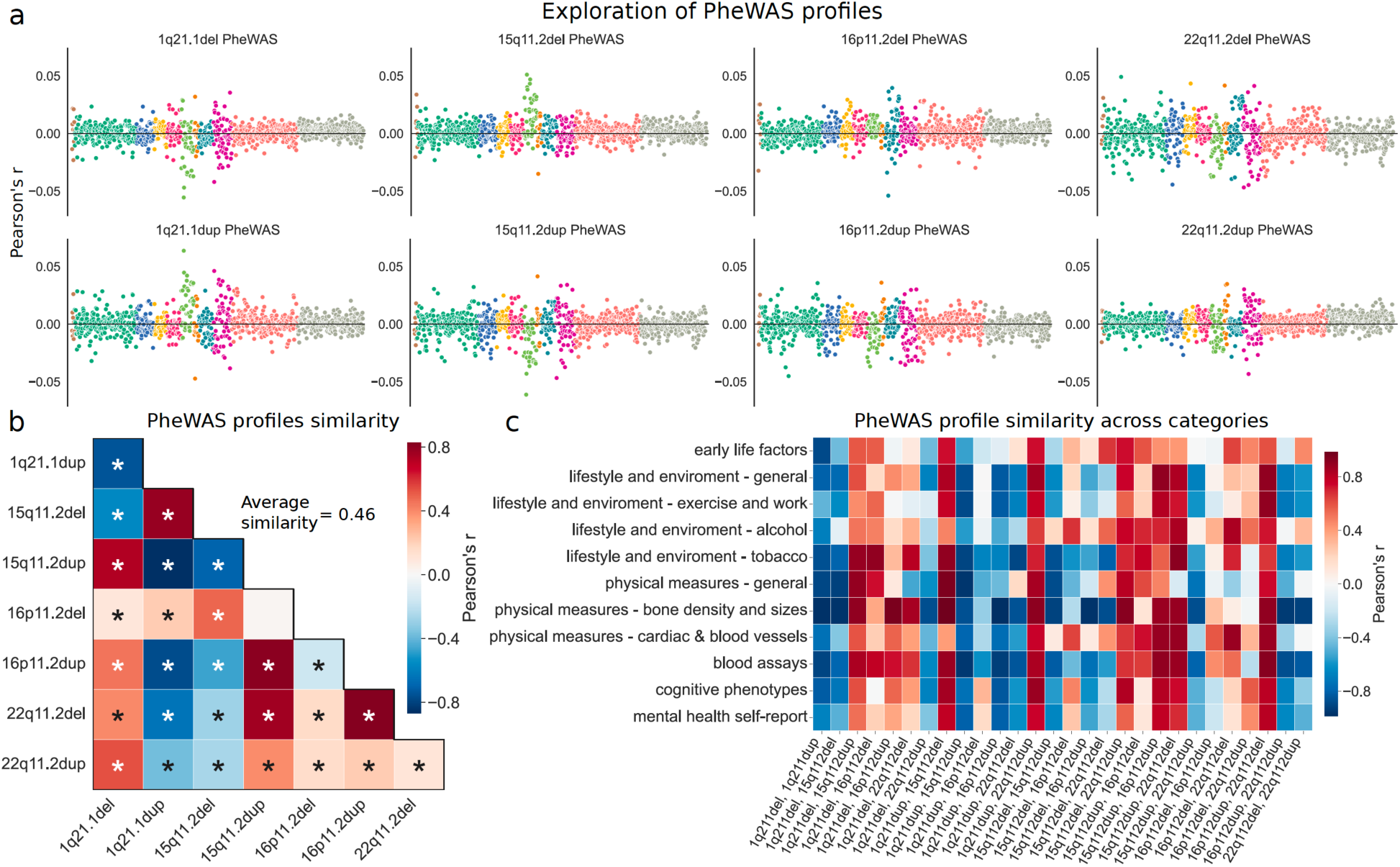
Eight different CNVs converge on similar phenome-wide association profiles. We carried out the PheWAS analysis for each intermediate phenotype to quantify the differences and commonalities in phenotypical consequences due to the eight CNVs. a) Depicts the obtained Pearson’s correlations from PheWAS analysis for each CNV status. Among those, 22q11.2 deletion emerges to show the strongest associations with numerous phenotypes across categories. Colors indicate the eleven categories. b) Linear association strength between Pearson’s correlations from the PheWAS analysis across all CNVs. Strong correlations suggest that CNVs are linked with similar phenotypes. Average similarity exceeds those of volumetric Cohen’s *d* maps and intermediate phenotypes. Asterisk denotes FDR-corrected significant correlations. c) Linear association strength between category-specific Pearson’s correlations from the PheWAS analysis across all CNVs. Detailed visualization depicts the similarity of the impact of CNVs on all phenotype categories. The direction of the linear relationship tends to be identical across categories for a given CNV pair (strong negative or strong positive), unlike across CNV pairs for a given category. Although CNVs entailed different brain changes (cf. Fig. 1-3), they exhibited similar PheWAS profiles, especially in categories such as bone density, blood assays, and general physical measures.

We subsequently zoomed in on the strong convergence across the phenotypic profiles characterizing each CNV by computing the correlation between CNV-phenotypic associations within each of the eleven considered categories (Fig. 5c). In particular, we found the bone density and sizes along with blood assays categories showed strong associations across CNV intermediate phenotypes, suggesting similar behavior within these categories. Altogether, the strong correspondences among CNV pairs suggest that CNV brain profiles are linked to similar phenotypes across all eleven categories.

### Detailing shared and distinct phenotypic associations

To shed light on which particular phenotypes are most strongly associated with CNV-specific brain signatures, we calculated the mean absolute Pearson’s correlations across the eight PheWAS analyses. Across all CNVs, diastolic blood pressure, alkaline phosphatase, and red blood cell count showed the strongest associations (Fig. 6a). Moreover, we examined which phenotypes are most consistently associated with CNV brain profiles. We found eight phenotypes associated with six CNV intermediate phenotypes and eleven phenotypes shared by five CNV intermediate phenotypes (Supp. Fig. 2b, c). The most consistently overlapping phenotype hits were from the blood assays category (e.g., mean corpuscular volume, SHBG, IGF-1), along with weight or home population density. In total, these robust and shared phenotypic associations point to the fact that CNV brain profiles are associated with similar systemic phenotypes.

**Figure 6.**
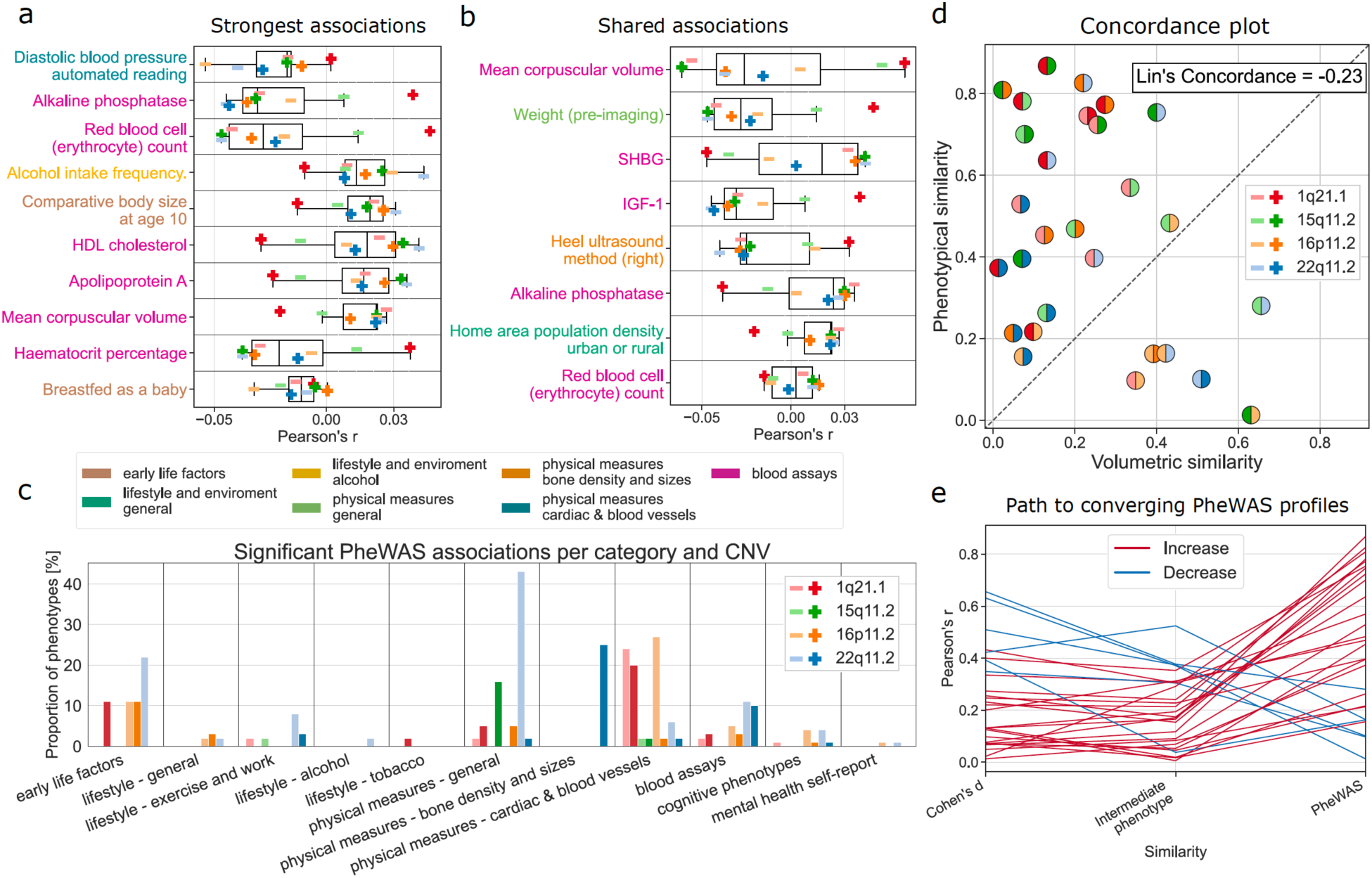
Detailing aspects convergence in phenome-wide portfolios across different CNVs. For all eight CNVs, we delineate the most prominent as well as distinctive associations among their PheWAS profiles in the UK Biobank. We also compare the different levels of gene-brain-behavior similarities. a) Phenotypes from the PheWAS analysis most strongly associated with the eight CNVs. We show ten phenotypes with the strongest average Pearson’s correlations across all CNVs. The most prominent association across CNVs is with diastolic blood pressure. b) Phenotypes most consistently associated with the eight CNVs. We find eight phenotypes that are associated with most (six) of the CNVs. Phenotypes are ordered according to the mean strength of the association. Most of the phenotypes are from the blood assays category. c) Number of significant hits per category for each intermediate phenotype conditioned on the shared phenotypical profile. For each of the eight intermediate phenotype expressions, we regressed out the remaining seven. Even after conditioning on the shared phenotypical associations, each particular CNV still shows a specific set of distinct phenome-wide associations across various categories. For example, 22q11.2 deletion still displays a high number of associations in physical measures - general category. d) Concordance between brain volume effects and PheWAS effects. The absolute value of correlation between Cohen’s *d* brain maps (Fig. 1a) is plotted against the absolute value of correlation between PheWAS profiles. Negative Lin’s concordance correlation hints at the disparity between volumetric and phenotypical similarity. Moreover, the majority of points lie above the 45°-degree line suggesting that PheWAS similarities are more substantial than volumetric similarities. e) From diverging brain patterns to converging portfolios. Each line represents an absolute Pearson’s correlation of Cohen’s *d* map, intermediate phenotype, and PheWAS profile for a given CNV pair. Convergence on PheWAS profiles is demonstrated by the increase in similarity in 22 of 28 CNV pairs. Hence, the similarity of CNV portfolios exceeded those of volumetric intermediate phenotypes.

Comparisons of the phenotypical profiles associated with each CNV intermediate phenotype revealed that though the phenotypical profiles are similar, there still remains unexplained residual variance as suggested by a maximum absolute association strength of r = 0.81. To access this remaining part of the variance, we computed new brain profiles adjusting for the other CNVs. Specifically, for each CNV-specific intermediate phenotype, we singled out the variation explained by the remaining seven. Thus, we obtained a set of eight unique intermediate phenotypes, each with the variation shared with other intermediate phenotypes removed. Subsequently, we used this new set to perform the PheWAS analysis and counted the relative number of associations surpassing the Bonferroni correction in each category. We still observed significant associations across CNVs and categories even after conditioning out on the shared associations. In particular, 22q11.2 deletion showed a high relative number of associations in the physical measures category (Fig. 6e). As such, CNVs did display unique phenotypic associations relative to other CNVs.

### Quantifying the path toward converging phenotypical profile

The observed magnitude of similarity between the phenotypic profiles of the CNV intermediate phenotypes reaching Pearson’s r = 0.84 demonstrated a strong relationship between phenotypic profiles across the 977 indicators. In general, the phenotypic similarity (absolute Pearson’s correlation of PheWAS outcomes) between CNVs exceeded their morphological similarity (absolute Pearson’s correlation between Cohen’s d maps) (Fig. 6d). The dissonance between the two similarity measures was highlighted by Lin’s concordance correlation coefficient equal to -0.23, suggesting poor concordance. More specifically, 22 of 28 CNV pairs showed stronger phenotypical similarity compared to volumetric similarity. Thus, CNVs were characteristic of stronger phenotypic signature associations in comparison to associations among volumetric signatures or intermediate phenotypes (Fig. 6e).

Our collective analyses demonstrated that although each CNV displays largely distinct whole-brain morphometric signatures, they converged on similar phenotypic profiles. In proving this, we transferred the intermediate phenotypes derived in the clinical cohort to the UK Biobank population cohort with 39,085 subjects. Using the subject-specific expression levels of eight intermediate phenotypes from eight rare CNVs allowed us to characterize complex phenotypical profiles of each CNV, providing a newly detailed portrait of their commonalities and idiosyncrasies.

## Discussion

CNVs offer a unique window of opportunity into the consequences of localized genetic variation on human traits. This is especially the case, given their known genetic architecture and typically high penetrance. In the present study, we built computational bridges between eight key CNVs in a multisite clinical dataset, on the one hand, and their deep phenotypic profiling in 39,085 subjects from the wider population, on the other hand. To this end, we designed an analytic framework that can quantitatively dissect the impact of distinct genetic mutations on brain organization and behavioral differentiation. Bringing over derived CNV-specific intermediate phenotypes to the population cohort revealed that the CNVs are tied to pleiotropic associations beyond physical and cognitive domains. This phenome-wide analysis across ∼1000 phenotypes revealed many ramifications for several body systems. Our collective analyses reveal wide-ranging similarities between the PheWAS profiles of the eight CNVs. Therefore, the phenotypic level appears to be the point of alignment for distinct long-segment genetic variants that we show to cause diverging morphological changes in brain morphology. Such late convergence in phenotypic consequences speaks to profound basic science questions regarding the organization of genetic influences on human brain and behavior.

For a long time, inquiries targeting genetic influences have been limited by the lack of longitudinal and deep multimodal measures of brain and behavior in large subject samples (Raznahan et al., 2022). Studies aimed at elucidating genotype-phenotype links were challenged by several challenges, including ascertainment bias, limited statistical power, and patchy phenotypic coverage (Modenato et al., 2021b). We are unlikely to have access to large enough clinical datasets soon – a condition *sine qua non* for definitive tests of phenotypic overlaps and differences between genetic variants. To overcome several of these hurdles, we here put forward solutions that take advantage of intermediate CNV phenotypes, a term coined in research on psychiatric disorders (Gottesman and Gould, 2003). These refer to biological traits that lie in between an individual’s external phenotype and innate genetic blueprint (Mark and Toulopoulou, 2016; Meyer-Lindenberg and Weinberger, 2006). As a core contribution of our present work, we demonstrate the added value of how intermediate phenotypes can be transferred for direct usage in other small hand-selected clinical cohorts and large-scale populational datasets. We captured CNV-specific intermediate phenotype representations as “genetics-first” whole-brain signatures derived from our clinical boutique dataset. By transferring these brain-wide representations over to the UK Biobank and carrying out PheWAS, we obtained systemic phenotypic associations across eleven rich phenotypic categories. Moreover, the PheWAS associations of the intermediate CNV phenotypes were concordant with previous studies investigating more circumscribed links between CNV status and indicators of cognitive performance, including fluid intelligence score (Kendall et al., 2019), physical measurements like weight or height (Auwerx et al., 2022; Owen et al., 2018), common medical conditions like hypertension or obesity (Crawford et al., 2019), and blood biomarkers like indicators of cholesterol fat metabolism pathways (Bracher-Smith et al., 2019). Our results highlight the potential of using intermediate phenotypes as a device to study a wide variety of rare conditions and thus accelerate the pace of neurogenetic innovation.

In addition to the difficulties in recruiting large high-quality clinical cohorts, today’s CNV research has been mainly restricted to child and adolescent patients referred to genetic clinics (Chawner et al., 2021). A narrow focus on younger populations favors finding deleterious effects on cognitive and behavioral phenotypes leading to recording high rates of developmental delay and early-onset medical conditions (Chawner et al., 2019; Crawford et al., 2019). However, by combining hand-crafted analytic solutions with recently emerged data resources, we here deliver evidence of pleiotropic consequences of CNVs invoking data from the general population. These consequences include systemic associations outside the central nervous system. This underappreciated insight is reflected in our results, including strong brain-behavior associations of the CNV profile in the UK Biobank population with blood pressure, cholesterol, or height and weight. Since CNVs do not show complete penetrance in all cases (Kirov et al., 2014), such associations portray a necessary picture of a broad spectrum of outcomes later in life. Hence, the constellation of results advocates rebalancing the medical care of CNV carriers towards more comprehensive medical monitoring in a broader patient pool (Chawner et al., 2021).

Our data-driven findings advocate a revised view of CNV pathophysiology as contributing to systemic illness. In a similar way, previous clinical research has provided evidence that schizophrenia and related psychotic disorders often affect multiple body systems (e.g., nervous, immune, or endocrine), even from illness onset (Alessi and Bennett, 2020; Guest, 2017). Pillinger and colleagues (2019) reported robust alterations in immune and cardiometabolic systems of a magnitude comparable to alterations in the central nervous system. Further examples of major brain disorders accompanied by problems outside the brain include gastrointestinal disorders in autism (Kohane et al., 2012), loss of bone density in depression (Sotelo and Nemeroff, 2017), or cardiovascular symptoms in bipolar disorder (Leboyer et al., 2012). Finally, a recent study showed that genetic liabilities for five major psychiatric disorders are associated with long-term outcomes in adult life, including socio-demographic factors and physical health (Leppert et al., 2020). Our findings thus add new pieces of knowledge that illuminate how the nervous system is interlocked with the rest of the body in a way that affects general well-being.

In concordance with reported system-wide effects in major brain disorders, our computational assays lay out pleiotropic associations in CNV carriers that go beyond mere cognitive domains. As one of many examples, we demonstrated how intermediate phenotypes tied to 22q11.2 deletion relate to an array of phenotypes in blood assays as well as cardiac and blood vessels categories. Moreover, similar to Auwerx and colleagues (2022), six of our eight examined CNVs were associated with body weight, insulin-like growth factor 1, alkaline phosphatase, or mean red blood cell volume. Therefore, both for CNVs and brain disorders, the associated bodily alterations may not be mere secondary effects (Pillinger et al., 2019). Instead, systemic manifestations could be a fundamental aspect of the primary biology of brain disorders. Critically, they might also lead to a reduced life span, as suggested by the 63% probability of survival to age 50 in adult carriers of 22q11.2 deletion (Crawford et al., 2019; Van et al., 2019). Moreover, psychotic disorders have been linked with 15–20 years shorter life expectancy (Kessler et al., 2007). Most of this premature mortality is predominantly due to elevated cardiovascular risk factors (Correll et al., 2017; Hoang et al., 2013) – causes that belong to the phenotype category among the most consistent associations in our phenome-wide assays. Thus, our findings speak in favor of CNVs as a complex disorder with several manifestations outside the brain that have considerable deleterious impacts on various parts of everyday lives.

More broadly, understanding pathophysiological disease mechanisms will be propelled by further disentangling the perplexing link between genes, brain, and behavior (Thompson et al., 2001). There is an active debate on the extent to which distinct gene dosage disorders can lead to different non-overlapping phenotypical profiles (Raznahan et al., 2022). This discourse was sparked from the observations that many or most SNPs and CNVs increase the risk for schizophrenia or autism (Bacchelli et al., 2020; Marshall et al., 2017). Polygenicity and pleiotropy, key features of the genetics underpinning psychiatric disorders (Gratten and Visscher, 2016; Moreno-De-Luca and Martin, 2021), imply that genetic mutations can converge on shared mechanisms at some level of pathway cascades, from genes to large-scale brain networks to the phenome. Here, we report a low similarity of intermediate phenotypes representing morphological CNV-specific brain signatures, in line with a documented broad diversity of regional morphometry patterns across genomic loci (Modenato et al., 2021b; Moreau et al., 2021a; Seidlitz et al., 2020). Conversely, the ramifications of carrying distinct CNV variants for cognition and behavior have previously been hypothesized to be more similar than those on brain anatomy (Raznahan et al., 2022; Silva et al., 2022). We here find strong evidence for substantial convergence of phenotypic measures across CNVs quantified by increased phenotypical similarity. Specifically, we observed a high degree of similarity between the phenotypical profiles (mean similarity r = 0.46 as measured by Pearson’s correlation across the CNV’s corresponding PheWAS profiles), which largely exceeded the similarity of brain morphometry profiles (mean similarity r = 0.2 as measured by the correlation of volumetric Cohen’s *d* maps). Based on the presented strong resemblance of phenotypic profiles of the examined eight CNVs, we speculate that the polygenic architecture of human phenotypic traits may be related to genotype-phenotype convergence that occurs later than on molecular pathways or macroscopic brain networks.

In conclusion, we have triangulated i) a purpose-designed analytical strategy, ii) a roadmap for investigating rare brain pathologies employing intermediate phenotypes derived from smaller clinical datasets and iii) a framework for application in population-scale cohorts. Our collective findings have far-reaching implications for the theoretical and empirical understanding of genotype-phenotype correspondences. Specifically, the observed phenotypic convergence sheds new light on why so many CNVs increase the risk for the same developmental, psychiatric disorders. Finally, building on our results, future studies may contextualize observed phenotypic convergences with respect to the known genetic architecture of CNVs to elucidate principles of brain development in health and disease.

## Online Methods

### Multisite clinical cohort

Our clinical dataset consisted of volumetric measurements derived from magnetic resonance imaging (MRI) brain scans from 860 subjects: 548 CNV carriers and 312 controls not carrying any CNV (Tab. 1). An extensive description of methods and analyses is available in an already published study with an identical dataset (Modenato et al., 2021a). In short, PennCNV and QuantiSNP were used, with standard quality control metrics, to identify CNVs. Structural brain-imaging data of CNV carriers were selected based on the following breakpoints according to the reference genome GRCh37/hg19: 16p11.2 proximal (BP4-5, 29.6-30.2MB), 1q21.1 (Class I, 146.4-147.5MB & II, 145.3-147.5MB), 22q11.2 (BPA-D, 18.8-21.7MB) and 15q11.2 (BP1-2, 22.8–23.0MB). Control individuals did not carry any CNV at these loci. Signed consents were obtained from all participants or legal representatives prior to the investigation. The CNV carriers were either probands referred to the genetic clinic for the investigation of neurodevelopmental and psychiatric disorders or their relatives (parents, siblings, and other relatives). 15q11.2del and 15q11.2dup CNVs were identified in the UK Biobank. Their respective carriers were added to our clinical cohort. Controls were either non-carriers within the same families or individuals from the general population. Furthermore, controls were carefully matched for sex and age to CNV carries.

### Clinical MRI data recording and processing

We analyzed a data sample of T1-weighted (T1w) images at 0.8–1 mm isotropic resolution. All T1w included in the analysis were quality checked by a domain expert (Modenato et al., 2021a). Data for Voxel-Based Morphometry were preprocessed and analyzed with SPM12 (http://www.fil.ion.ucl.ac.uk/spm/software/spm12/) (Ashburner, 2007; Ashburner and Friston, 2005; Lorio et al., 2016) running under MATLAB R2018b (https://www.mathworks.com/products/new_products/release2018b.html). Further quality control was performed using standardized ENIGMA quality control procedures (http://enigma.ini.usc.edu/protocols/imaging-protocols/). Finally, neurobiologically interpretable measures of gray matter volume were extracted in all participants by summarizing whole-brain MRI maps in the MNI reference space. This feature-generation step was guided by the topographical brain region definitions of the commonly used Schaefer-Yeo atlas with 400 parcels (Schaefer et al., 2018). The derived quantities of local gray matter volumetry resulted in 400 volume measures for each participant. The subject-level brain region volumes provided the input variables for our linear discriminant analysis (cf. below). As a data-cleaning step, derived regional brain volumes were adjusted for total grey matter, age, age^2^, and sex as fixed effects and scanning site as a random factor, following previous research on this dataset (Modenato et al., 2021a).

### Population data source

The UK Biobank (UKBB) is the largest biomedical resource that offers extensive behavioral and demographic assessments, medical and cognitive measures, as well as biological samples in a cohort of ∼500,000 participants recruited from across Great Britain (https://www.ukbiobank.ac.uk/). This openly accessible population dataset aims to provide brain-imaging for ∼100,000 individuals, planned for completion in 2023. The present study was based on the recent brain-imaging data release from February/March 2020. Concretely, our data sample included measurements from 39,085 participants with brain-imaging measures and expert-curated image-derived phenotypes of gray matter morphology (T1-weighted MRI). Among the participants, 48% were men and were 52% women with age between 40 and 69 y.o. when recruited [mean age 55 y.o., standard deviation (SD) 7.5 y.]). We benefited from the uniform data preprocessing pipelines designed and implemented by the FMRIB, Oxford University, Oxford, UK (Alfaro-Almagro et al., 2018) to improve comparability and reproducibility. The present analyses were conducted under UK Biobank application number 25163. All participants provided written informed consent. For further information on the consent procedure, refer to http://biobank.ctsu.ox.ac.uk/crystal/field.cgi?id=200.

MRI scanners (3T Siemens Skyra) at several dedicated data collection sites used matching acquisition protocols and standard Siemens 32-channel radiofrequency receiver head coils. To protect the anonymity of the study participants, brain-imaging measures were defaced, and any sensitive meta-information was removed. Automated processing and quality control pipelines were deployed (Alfaro-Almagro et al., 2018; Miller et al., 2016). To improve the homogeneity of the brain-imaging scans, the noise was removed using 190 sensitivity features. This approach allowed for the reliable identification and exclusion of problematic brain scans, such as due to excessive head motion.

The structural MRI data were acquired as high-resolution T1-weighted images of brain anatomy using a 3D MPRAGE sequence at 1mm isotropic resolution. Preprocessing included gradient distortion correction, the field of view reduction using the Brain Extraction Tool (Smith, 2002) and FLIRT (Jenkinson et al., 2002), as well as non-linear registration to MNI152 standard space at 1 mm resolution using FNIRT (Andersson et al., 2007). To avoid unnecessary interpolation, all image transformations were estimated, combined, and applied by a single interpolation step. Tissue-type segmentation into the cerebrospinal fluid, grey matter and white matter to generate full bias-field-corrected images was achieved using FAST (FMRIB’s Automated Segmentation Tool, (Zhang et al., 2001)). Finally, grey matter images were used to extract gray matter volumes in parcels according to the Schaefer-Yeo atlas with 400 regions (Schaefer et al., 2018). Following previous work on the UKBB (Schurz et al., 2021; Spreng et al., 2020), inter-individual variations in brain region volumes that could be explained by nuisance variables of no interest were adjusted for by regressing out: body mass index, head size, head motion during task-related brain scans, head motion during resting-state fMRI scanning, head position and receiver coil in the scanner (x, y, and z), position of scanner table, as well as the data acquisition site.

### Statistical analysis for volumetric brain measures

All subsequent analyses were performed in Python v3.8 as scientific computing engine (https://www.python.org/downloads/release/python-380/). We used Cohen’s *d* to quantify the effect size of the CNVs on individual regional volumes d (Cohen, 2013). For a given region, Cohen’s *d* is defined as:

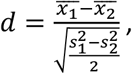

where 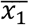 corresponds to the mean region volume across CNV carriers, 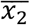 corresponds to the mean region volume across controls. Similarly, *s*_1_ and *s*_2_ correspond to standard deviations of CNV carriers and controls.

We compared Cohen’s *d* volumetric brain maps (and intermediate phenotypes brain maps) between different CNVs using Pearson’s correlation. Furthermore, we used spin permutation testing to calculate empirical p-values for the ensuing correlation coefficient (Alexander-Bloch et al., 2013; Reardon et al., 2018).

Finally, we calculated Lin’s concordance correlation coefficient to quantify the agreement of similarities between volumetric Cohen’s *d* maps, intermediate phenotypes, and PheWAS profiles (Lawrence and Lin, 1989). The degree of concordance between the two measures is thus calculated as:

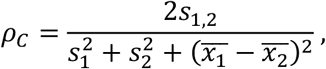

where *s*_1,2_ corresponds to covariance between *x*_1_ and *x*_2_.

### Charting complex association using phenom-wide association study

We performed a rich annotation of the derived intermediate phenotypes by means of a phenome-wide association analysis benefitting from a wide variety of almost 1,000 lifestyle factors. For a detailed description of phenotype extraction and analysis, refer to our previously published studies (Saltoun et al., 2022; Savignac et al., 2022). Feature extraction was carried out using two utilities designed to obtain, clean and normalize UKBB phenotype data according to predefined rules. In short, we collected a raw set of ∼15,000 phenotypes that we further processed by the FMRIB UKB Normalisation, Parsing And Cleaning Kit (FUNPACK version 2.5.0; https://zenodo.org/record/4762700#.YQrpui2caJ8). FUNPACK is designed to perform automatic refinement on the UKB data, which included removing ‘do not know’ responses and filling the blank left by unanswered sub-questions. The FUNPACK-derived phenotype information covered 11 major categories, including cognitive and physiological assessments, physical and mental health records, blood assays, as well as sociodemographic and lifestyle factors. The output consisted of a collection of 3,330 curated phenotypes which were then fed into PHEnome Scan ANalysis Tool (PHESANT (Millard et al., 2018), https://github.com/MRCIEU/PHESANT) for further refinement in an automated fashion. PHESANT performs further data cleaning and normalization along with labeling data as one of four data types: categorical ordered, categorical unordered, binary, and numerical. Categorical unordered variables were one-hot encoded, such that each possible response was represented by a binary column (true or false). The final curated inventory comprised 977 phenotypes spanning 11 FUNPACK-defined categories. Furthermore, we used Pearson’s correlation to quantify the association strength between these 977 phenotypes with subject-specific expressions of our eight intermediate phenotypes (cf. below). To ensure that the correlations are not driven by a few outlying intermediate phenotype expressions, we first discarded 551 subjects based on Tukey’s interquartile range rule for outlier detection (Hodge and Austin, 2004).

### Multi-class prediction model and intermediate phenotype extraction

Technically, our core aim was to derive robust CNV-specific representations of intermediate phenotypes from a clinical sample that could be transferred to a large population resource for deep profiling. We derived the intermediate phenotypes as systematic co-deviations of brain morphometry attributable to each of our eight target CNVs. To this end, we capitalized on linear discriminant analysis to extract separating rules between CNV carriers and controls based on whole-brain volume measurements. Linear discriminant analysis can be viewed as a generative approach to classifying CNV carriers, which requires fitting multivariate Gaussian distribution to regional brain volumes and produces a linear decision boundary (Hastie et al., 2009). Using a linear model represents a data-efficient and directly biologically interpretable approach to our analysis, especially in our boutique datasets with limited numbers of subject samples (Bzdok and Ioannidis, 2019). These datasets are characteristic of the low sample size regularly encountered in biology and medicine, which typically impedes the application of more complex non-linear models that require high numbers of parameters to be estimated (Bzdok and Yeo, 2017). In particular, LDA-derived discriminant vectors/functions served to represent CNV-specific intermediate phenotypes.

As another key model property of direct relevance to our present analysis goals, LDA can also be viewed as a dimensionality technique. This is because this modeling framework enables the extraction of underlying coherent principles among our anatomical target regions that are most informative in telling apart CNV carriers from controls. To do so, LDA has access to class labels (CNV status in our case) and thus belongs to supervised techniques (Hastie et al., 2009). Specifically, LDA projects the input subjects’ set of brain morphology measurements into a linear subspace, consisting of the directions which maximally separate our classes (Hart et al., 2000). This dimensionality reduction quality of LDA was a necessary prerequisite for the ability to extract intermediate phenotypes from one dataset and transfer them to other datasets.

In our study, we used LDA to mostly classify between CNV carriers and controls. Specifically, we derived a single LDA prototype for each CNV status to obtain eight CNV-specific intermediate phenotypes. We next introduce the algorithm on the example of 22q11.2 deletion and controls. The algorithm corresponds to Fisher’s implementation of the LDA model based on solving the eigenvalue decomposition problem. We first describe the algorithm for dimensionality reduction. The goal of our dimensionality reduction was to provide biologically interpretable low-rank views on the 22q11.2 deletion and controls in the form of a linear combination of brain region volumes. As a general rule, the maximum number of dimensions equals to the number of classes − 1. Therefore, since we here discriminated between two classes (controls and, say, 22q11.2 deletion), we obtained a one-dimensional vector that encapsulated the 22q11.2 deletion intermediate phenotype. In mathematical terms, we represented the set of *N* subjects (22q11.2 deletion carries plus controls) by a matrix *X* = {*x*_1,_ *x*_2_, …, *x*_*N*_ } where the vector *x*_1_ indicates subject 1 characterized by 400 brain volumes given by SchaeferYeo atlas. The transformation from the original brain atlas space to a low-rank embedding expression can be written as follows:

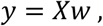

where *w* is the LDA derived discriminant function for how to project from the set of atlas regions to CNV distinction, *y* represents continuous CNV-control separation estimates as a vector of brain volumes projected into a one-dimensional subspace. The resulting vector *y* thus yielded a single compact estimate of the relative presence of the intermediate phenotype at hand for each subject.

Maximum CNV-control separation (i.e., the optimal subspace for classification) is achieved by maximizing the between-to-within group variability corresponding to finding a projection vector *w* that maximizes the ratio between-class scatter matrix and the within-class scatter matrix (Tharwat et al., 2017). The within-class scatter matrix *S*_*i*_ of a class *i* holds the class-specific distance between the subject’s regional volumes and the corresponding mean regional volumes:

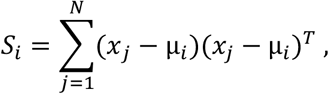

where *x*_*j*_ stands for the vector of 400 volumes of subject *j, N* is the number of subjects and µ_*i*_ is the mean of class *i*.

The total within-class scatter matrix *S*_*W*_ is calculated as the sum of within-class scatter matrices *S*_*i*_ across the two classes of controls and, say, 22q11.2 deletion:

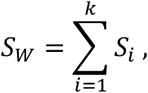

where *k* is the number of classes (*k* = 2 in our case).

In our two-class scenario, the between-class scatter matrix *S*_*B*_ represents the distance between the regional volume means across subjects of the two classes:

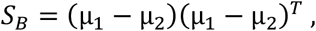

where µ_1_ correspond to mean region volumes of class 1.

In summary, Fisher’s between-to-within variance ratio represents an objective function for finding optimal decomposition of the classification subspace; here pertaining to identifying CNV carries. Concretely, Fisher’s criterion maximizes the ratio of the between-class variance *S*_*B*_ to the within-class variance *S*_*w*_ as a function of projection vector *w*:

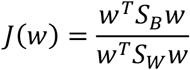

We can find the vector *w* if we re-write the ratio into a generalized eigenvalue-eigenvector problem:

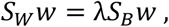

where λ represent the eigenvalues of the LDA projection vector *w*.

Then, the solution to this modelling problem and thus the LDA projection vector can be obtained by eigenvalue decomposition of 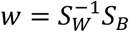.

LDA offers to go beyond mere case-control group differences by its natural capability for multi-class classification. We built a single LDA model to delineate differences among the eight CNVs by modelling each CNV as a separate class in single model estimation. In this scenario, the between-class scatter matrix represents the sum of distances between each CNV-specific mean regional volumes and the total mean regional volumes:

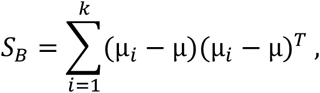

where µ represents mean region volumes across all subjects and classes.

Note that in multi-class classification projection vector *w* and projected values *y* can be extended to a multi-dimensional matrices *W* and *Y*. We projected 400 regional volumes to two dimensions that best discriminate between the eight individual CNV types.

Finally, the quantity µ_*k*_*ww*^*T*^ corresponds to CNV-specific LDA coefficients. In a two-class scenario, coefficients further reduce to (µ_1_ − µ_2_)*ww*^*T*^. These coefficients reveal the unique contribution of each brain region volume towards the separability of the CNV carries based on brain-wide morphology patterns. Therefore, the coefficients provide an understanding of the relative importance of the brain regions. Moreover, the LDA coefficients are estimated in a mutually dependent fashion. In other words, LDA coefficients are estimated hand-in-hand in conjunction with the other brain region volumes.

In order to find a low-rank subspace that maximally separates 22q11.2 deletion carriers and controls, we used the LDA projection vector *w* to re-express (i.e., more formally, project) the set of 400 regional volumes of a given subject onto a single dimension representing 22q11.2 deletion intermediate expression level signature. Moreover, as a step from dimensionality reduction to classification, projected subject morphologies can be used to construct a discriminant function. This function, used to assign class labels of 22q11.2 deletion or control, is generated by fitting class conditional densities *P*(*x*|*y* = *k*) to the low-dimensional data *x* of each class *k*. Conditional class probabilities are then obtained by using Bayes’ rule:

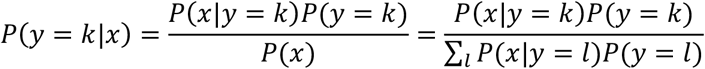

We need to calculate the class posteriors for optimal decision making. The decision is based on the selection of class *k*, which maximizes this posterior probability *P*(*y* = *k*|*x*). We model our two classes as two multivariate Gaussian distribution densities with a common covariance matrix. Therefore, we can re-write the class conditional distribution *P*(*x*|*y* = *k*) as:

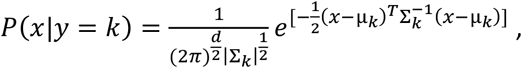

where *d* is the number of features (in our case 400 corresponding to SchaeferYeo regions), |Σ_*k*_| is the determinant of the covariance matrix of class *k*.

Assuming that each class has the same covariance matrix Σ_*k*_ = Σ (as in the present investigation), we can then simplify the posterior to:

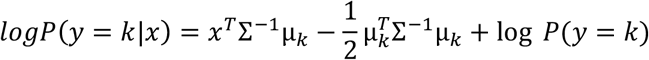

Finally, a participant’s brain volume features *x* is assigned to class *k* with the maximal posterior probability. As a consequence of this formula/model specification, LDA produces a linear decision boundary since the decision rule depends on *x* only through a linear combination of its elements.

### Building and validating robust prediction models

To recapitulate, our goal was to derive eight CNV-specific intermediate phenotypes using LDA. Therefore, we built separate CNV-specific LDA models designated to learn predictive principles to tell apart between CNV carriers and controls. However, we faced the challenge of the low number of CNV carriers. This challenge is inherent to various boutique datasets of rare medical conditions. Consequently, our number of measured features (regional brain volumes) was higher than the number of observation samples (subjects). Concretely, we disposed on average of 67 subjects per CNV class (cf. Table 1), while each subject was described by 400 regional volumes. Such a high-dimensional data scenario can lead to overfitting (Bzdok et al., 2019), where the model learns the detail and noise in the training samples and performs poorly in group classification on unseen test samples (Hastie et al., 2009). Hence, we used bootstrap aggregation (bagging), an ensemble learning method that can be used to reduce overfitting (Breiman, 1996). Bagging gains its value by profiting from a wisdom-of-crowds strategy. Concretely, we used a set of trained LDA models to obtain more robust and better predictive performance than could be obtained from a single trained LDA model in isolation (Breiman, 1996). Such a model-averaging design improves classification performance by reducing variance (Hastie et al., 2009).

**Table 1.** Clinical dataset demographics. CNV loci chromosome coordinates are provided with the number of genes encompassed in each CNV and with a well-known gene for each locus, to help recognize the CNV. Abbreviations, Del: deletion; Dup: duplication; ASD: autism spectrum disorder; SZ: schizophrenia; chr: chromosome; Age: mean age; SD: standard deviation; nGenes: number of genes.

| Loci | Chr (hg19)<br>start-stop | nGenes<br>(Gene) | Type | Subjects | Age<br>mean (SD) | Sex<br>Male/Female | Diagnosis<br>ASD/SZ |
| --- | --- | --- | --- | --- | --- | --- | --- |
| 1q21.1 | chr1<br>146.53-147.39 | 7<br>CHDIL | Del | 24 | 29 (18) | 9/15 | 1 |
|  |  |  | Dup | 15 | 34 (17) | 7/8 | 1 |
| 15q11.2 | chr15<br>22.81-23.09 | 4<br>CYFIP1 | Del | 112 | 55 (7) | 51/61 | 0 |
|  |  |  | Dup | 146 | 54 (7) | 69/77 | 0 |
| 16p11.2 | chr16<br>29.65-30.20 | 27<br>KCTD13 | Del | 80 | 17 (12) | 46/34 | 13 |
|  |  |  | Dup | 69 | 31 (14) | 37/32 | 11 |
| 22q11.2 | chr22<br>19.04-21.47 | 49<br>AIFM3 | Del | 69 | 17 (9) | 33/36 | 11 |
|  |  |  | Dup | 19 | 19 (14) | 12/7 | 2 |
| Controls |  |  |  | 312 | 26 (14) | 179/133 | 1 |

Therefore, we performed bagging during the derivation of LDA models separately for all eight CNV classes. Specifically, we used the following analytical strategy for a set of subjects consisting of a single CNV type and controls. In the first phase, a randomly perturbed version of the dataset is created by sampling the subject cohort with replacement. This bootstrap resampling served as the “in-the-bag” set of samples (i.e., subjects). The number of “in-the-bag” CNV carriers and controls equals their number in the dataset. Furthermore, the LDA model was trained on this training “in-the-bag” dataset. Model performance was then evaluated on all subjects from the dataset that were not selected for the “in-the-bag” dataset. These subject samples formed a testing “out-of-bag” dataset. The performance (i.e., classification accuracy) was based on the Mathews correlation coefficient, which has been reported to produce a more informative and truthful score than accuracy and F1 score (Chicco and Jurman, 2020).

We repeated the bootstrap resampling procedure with 100 iterations. In so doing, we obtained different realizations of the entire analysis process and ensuing LDA model estimate. Concretely, the bagging algorithm resulted in 100 trained LDA models that were used to obtain 100 out-of-bag predictions in unseen subjects. We calculated the final prediction accuracy as a mean across the 100 performance estimates. Critically, the average over the collection of separately estimated LDA discriminant functions served as our CNV-specific intermediate phenotype that provided the basis for downstream analysis steps. Finally, we characterized each subject by the intermediate phenotype expression level, which we calculated as the average one-dimensional LDA projection of regional volume sets across the 100 replications. In summary, the variance of local information in the 100 redraws of our original clinical subject cohort promoted diversity among the obtained candidate predictive rules, thus strengthening the fidelity of our ultimate predictions (Kuncheva and Whitaker, 2003).

To further safeguard against the risk of overfitting, we optimized the shrinkage parameter of each LDA model. Shrinkage corresponds to regularization used to stabilize the estimation of covariance matrices during model training. The empirical sample covariance is a poor estimator when the number of samples is small compared to the number of features (Chen et al., 2011). In particular, our sample covariance matrix contained 80,200 unique elements, which is almost 1200 times more than the average number of CNV carriers. Therefore, shrinkage helped improve the generalization performance of the LDA classifier. Using a nested cross-validation architecture, we performed a rigorous search over 11 shrinkage hyper-parameter choices between 0 and 1, in steps of 0.1, in each “in-the-bag” bootstrap iteration (GridSearchCV function from *sklearn*). The optimal hyperparameter choice was based on a leave-one-out strategy. In this cross-validation technique, each sample of the “in-the-bag” dataset was used once as a test set of unseen subjects, while the remaining subject samples formed the training set.

Finally, we evaluated the significance of a cross-validated score and thus assessed whether our ensemble LDA model displayed above-chance classification performance. Specifically, we carried out a label permutation test to quantify whether our LDA model outperforms the empirical null model. The null distribution was generated by calculating the prediction accuracy of our LDA classifier on 100 different permutations of the dataset. In these, features remained unchanged, but class labels (i.e., CNV carriers or controls) were randomly shuffled. Such a shuffling corresponded to the null hypothesis, which states there is no dependency between the features and labels. LDA model displayed above-chance classification performance if its prediction accuracy was higher than 97.5th percentile of prediction accuracy coefficient distribution derived from 100 permuted models.

### Performing model inspection using feature importance

After deriving robust LDA classifiers, we inspected which brain regions were the most informative in telling apart CNV carriers and controls. In other words, we aimed to contextualize and unpack the prediction rules of our ensemble LDA model. However, the bagging algorithm led to obtaining a collection of LDA models, resulting in a collection of estimates for each LDA coefficient and subject-specific intermediate phenotype expressions. Since each LDA model is trained on a different bootstrap population, it might happen that two distinct LDA models’ coefficients would carry opposite signs due to sign invariance of LDA dimensionality reduction. Therefore, we aligned all LDA models by multiplying them with -1 or 1 to produce a positive correlation between LDA coefficients and a corresponding Cohen’s d map.

Furthermore, we designed a criterion to test which LDA coefficients are significant, meaning which features significantly contribute to the classification. Significant coefficients had the distribution of 100 LDA coefficients significantly different from 0. Concretely, they were robustly different from zero if their two-sided confidence interval according to the 2.5/97.5% bootstrap-derived distribution did not include zero.

### Carrying intermediate phenotype expressions over for deep characterization in other data resources

The core contribution of our work lies in the study of derived intermediate phenotypes in the population dataset. To do so, we transferred the CNV-specific intermediate phenotypes carefully derived in our boutique dataset and quantified their expression in the general population (i.e., UK Biobank). Notably, UK Biobank itself contains CNV carriers. Therefore, we aimed to validate the transferability of intermediate phenotypes by testing the difference in intermediate phenotype expression between CNV carriers and controls in both the clinical dataset and UK Biobank. Specifically, we tested the null hypothesis of no difference in mean expression of intermediate phenotype in CNV carriers and controls. We adopted a two-sample bootstrap hypothesis test for means difference (Efron and Tibshirani, 1994). First, we shifted both expression levels to have the same mean. We then created bootstrap samples by drawing subjects with replication out of the shifted arrays. We computed the difference in means between the bootstrap distribution of intermediate phenotype expression levels. Finally, we constructed 1,000 of these bootstrap replicates. The hypothesis p-value corresponded to the fraction of replicates with a difference in means greater than or equal to what was observed.

