## Supplementary material for "Rare CNVs and phenome-wide profiling: a tale of brain-structural divergence and phenotypical convergence": Supp. Fig.

### Supplementary Figures

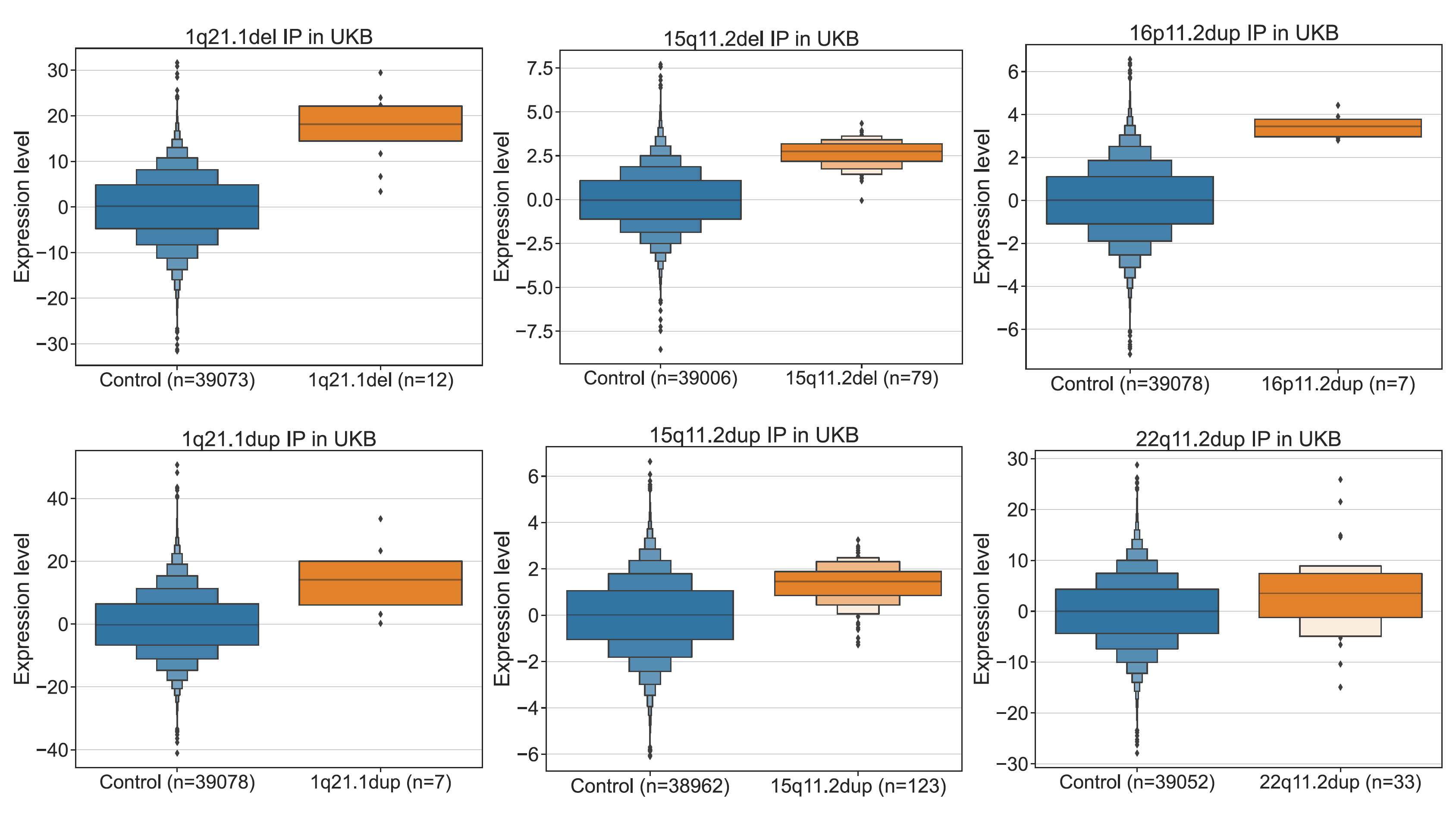

**Supplementary Figure 1**

**Intermediate phenotype expressions in the UK Biobank**

We quantified the presence of each CNV-specific intermediate phenotype derived from the clinical dataset. Here, we plot letter-value (boxen) plots for CNVs with at least five carriers in the UK Biobank. In all of these CNVs, the expression of respective intermediate phenotype was higher in actual CNV carries than in non-CNV carries.

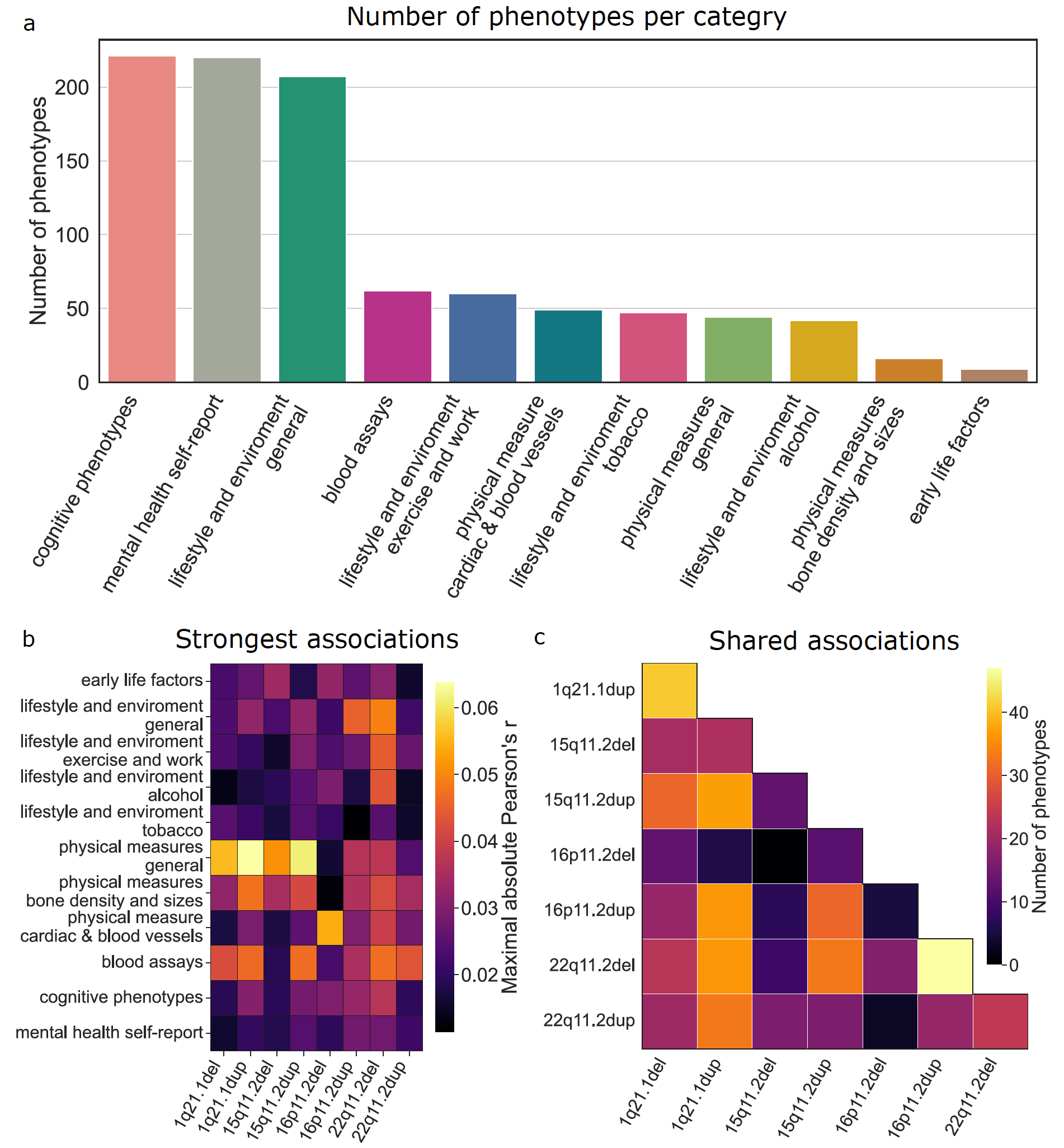

**Supplementary Figure 2**

**Dissecting PheWAS analyses across 11 categories**

We inspected which of our eleven categories are most strongly and most consistently associated with CNV status. a) The 977 available phenotypes span eleven categories. Three categories dominated our PheWAS, with more than 200 phenotypes per category. b) Strongest associations across categories and CNVs. We plot the maximal absolute Pearson’s correlation for each category and CNV. The strongest associations are in the physical measures – general and blood assays category. c) Shared associations across CNVs. The number of shared phenotypical associations varied between 0 and 43 (16p11.2 proximal duplication and 22q11.2 deletion). Moreover, the number of shared associations is significantly correlated with the phenotypical similarity between CNVs (r = 0.80, p < 10^-4^). These results support the high phenotypical similarities among different CNV loci.

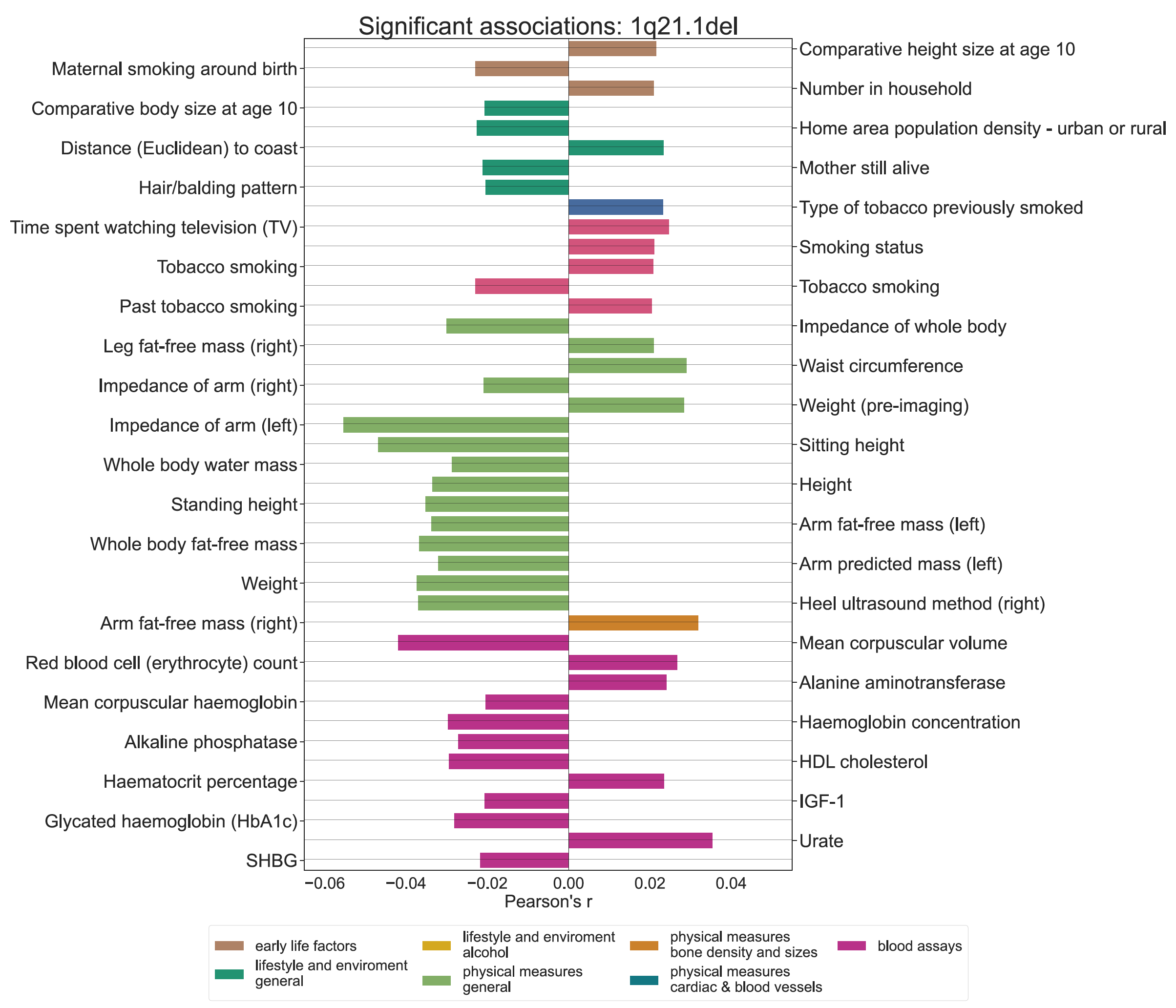

**Supplementary Figure 3**

Significant PheWAS associations for 1q21.1 distal deletion intermediate phenotype expression.

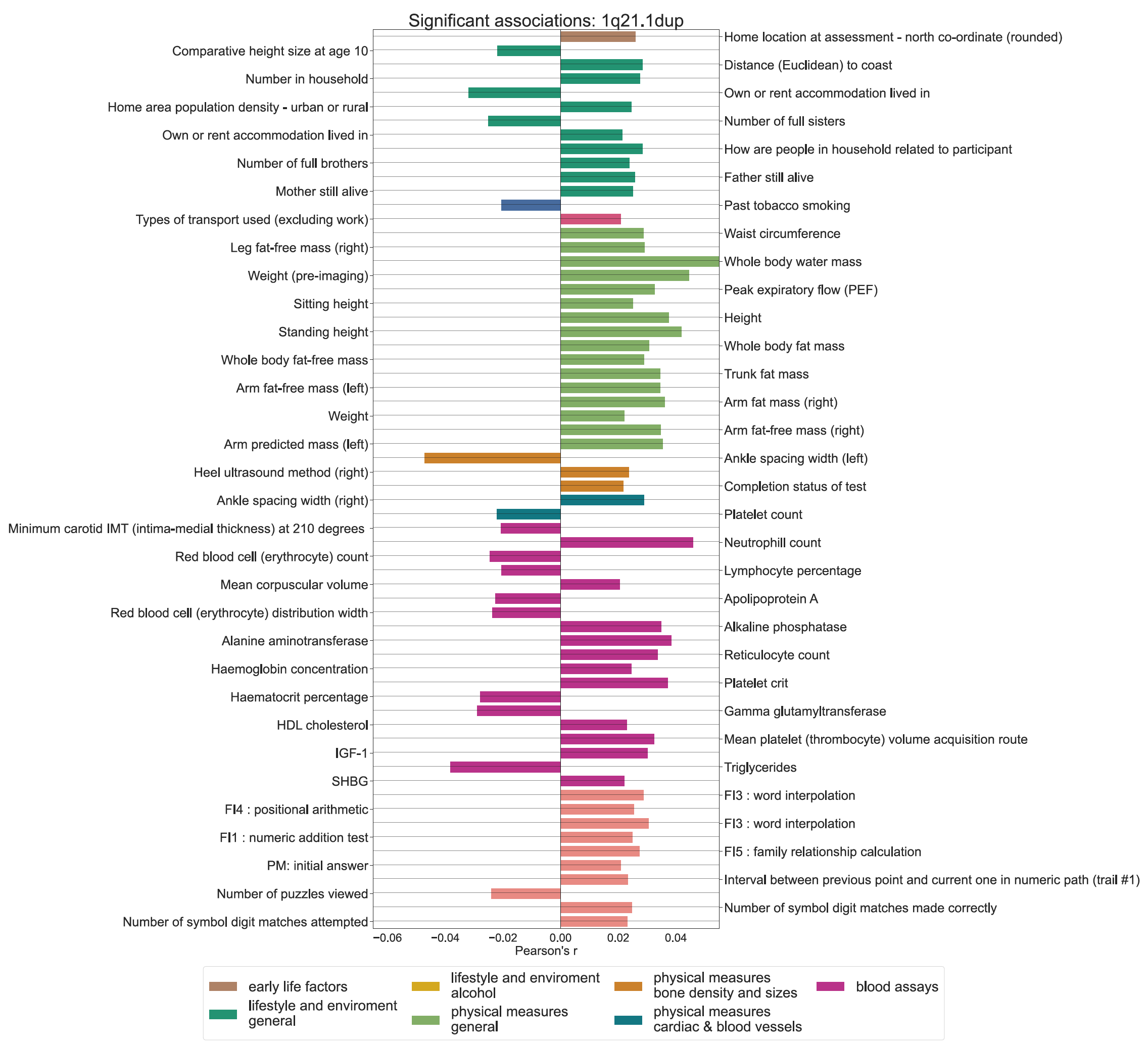

**Supplementary Figure 4**

Significant PheWAS associations for 1q21.1 distal duplication intermediate phenotype expression.

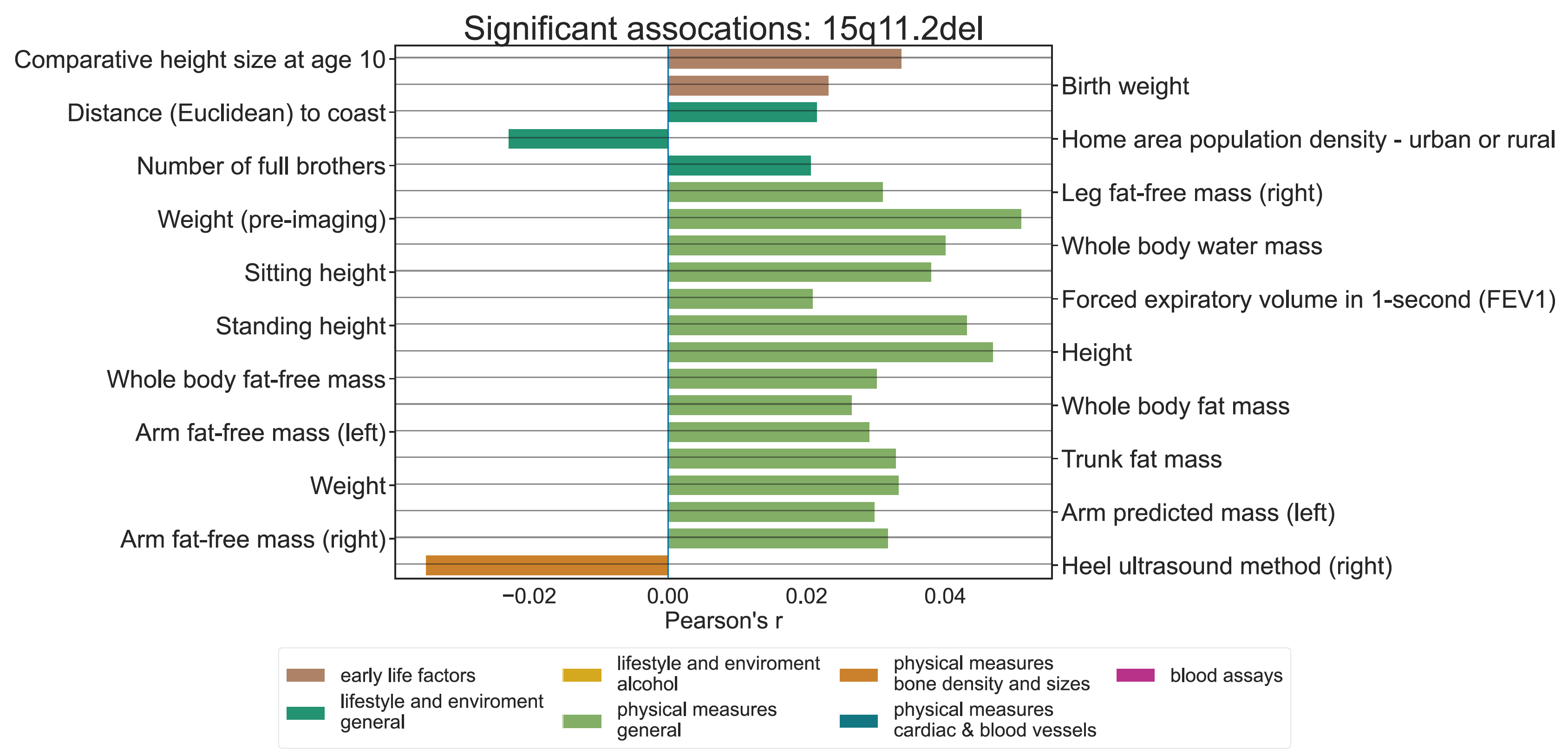

**Supplementary Figure 5**

Significant PheWAS associations for 15q11.2 deletion intermediate phenotype expression.

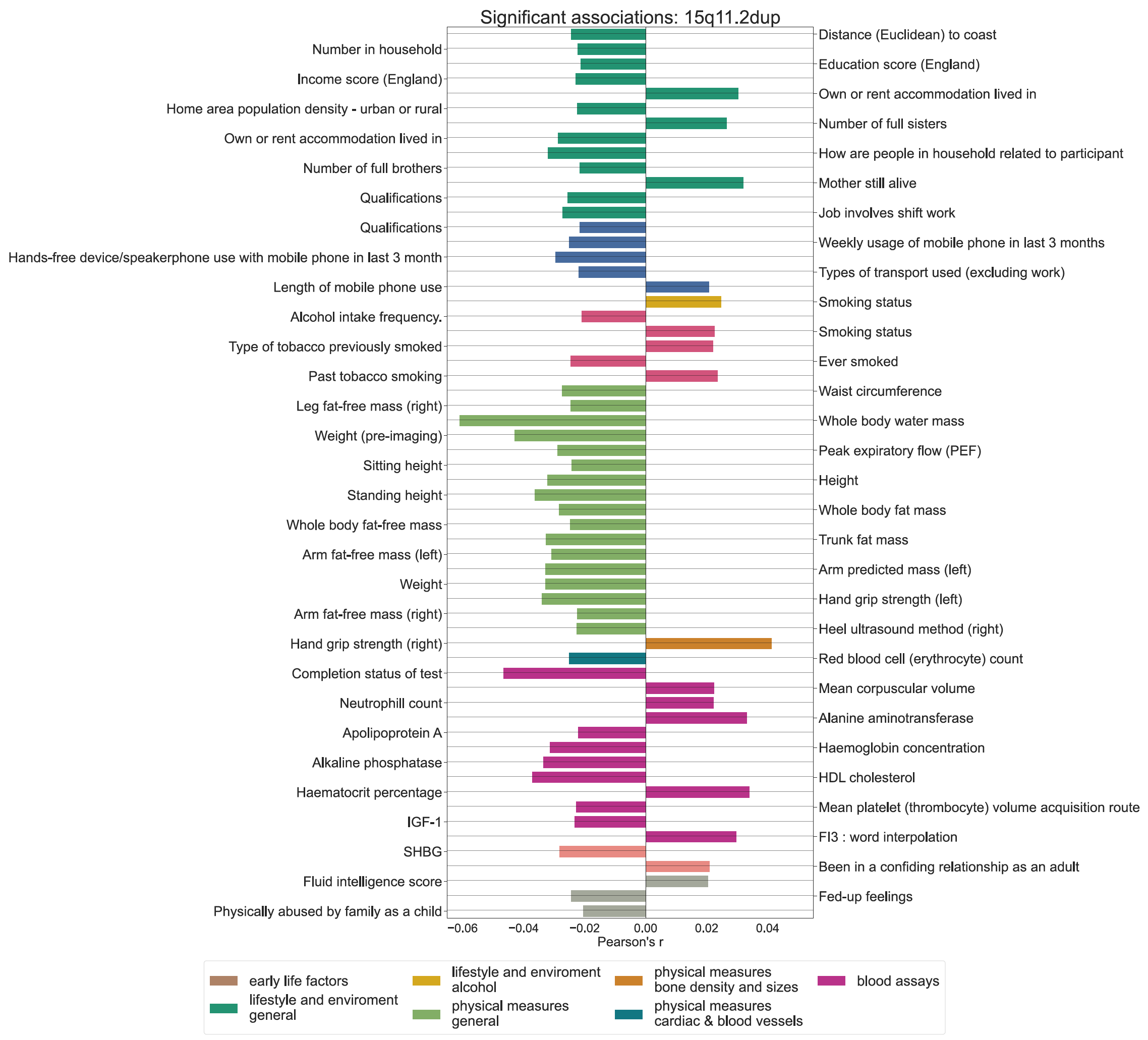

**Supplementary Figure 6**

Significant PheWAS associations for 15q11.2 duplication intermediate phenotype expression.

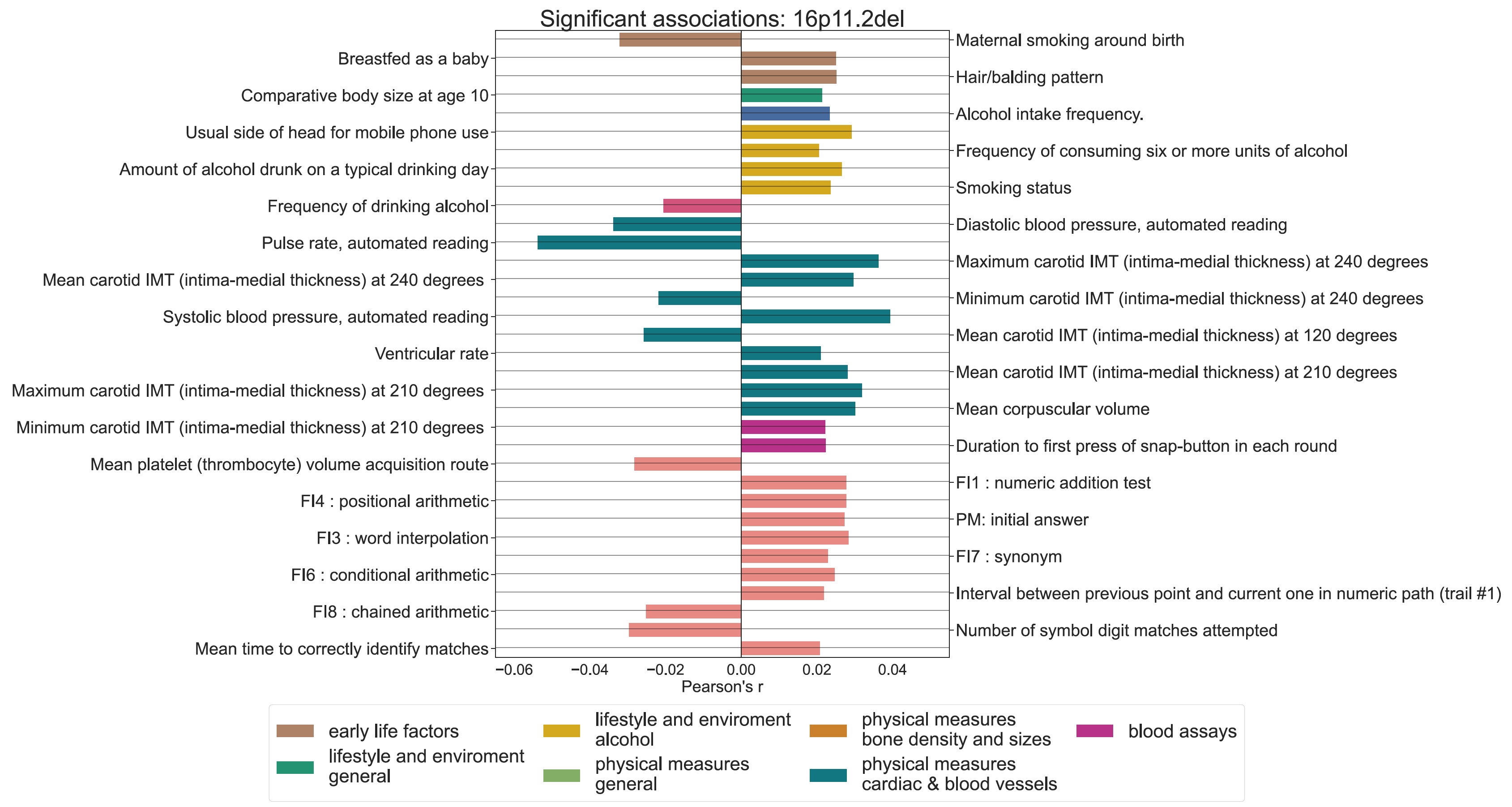

**Supplementary Figure 7**

Significant PheWAS associations for 16p11.2 proximal deletion intermediate phenotype expression.

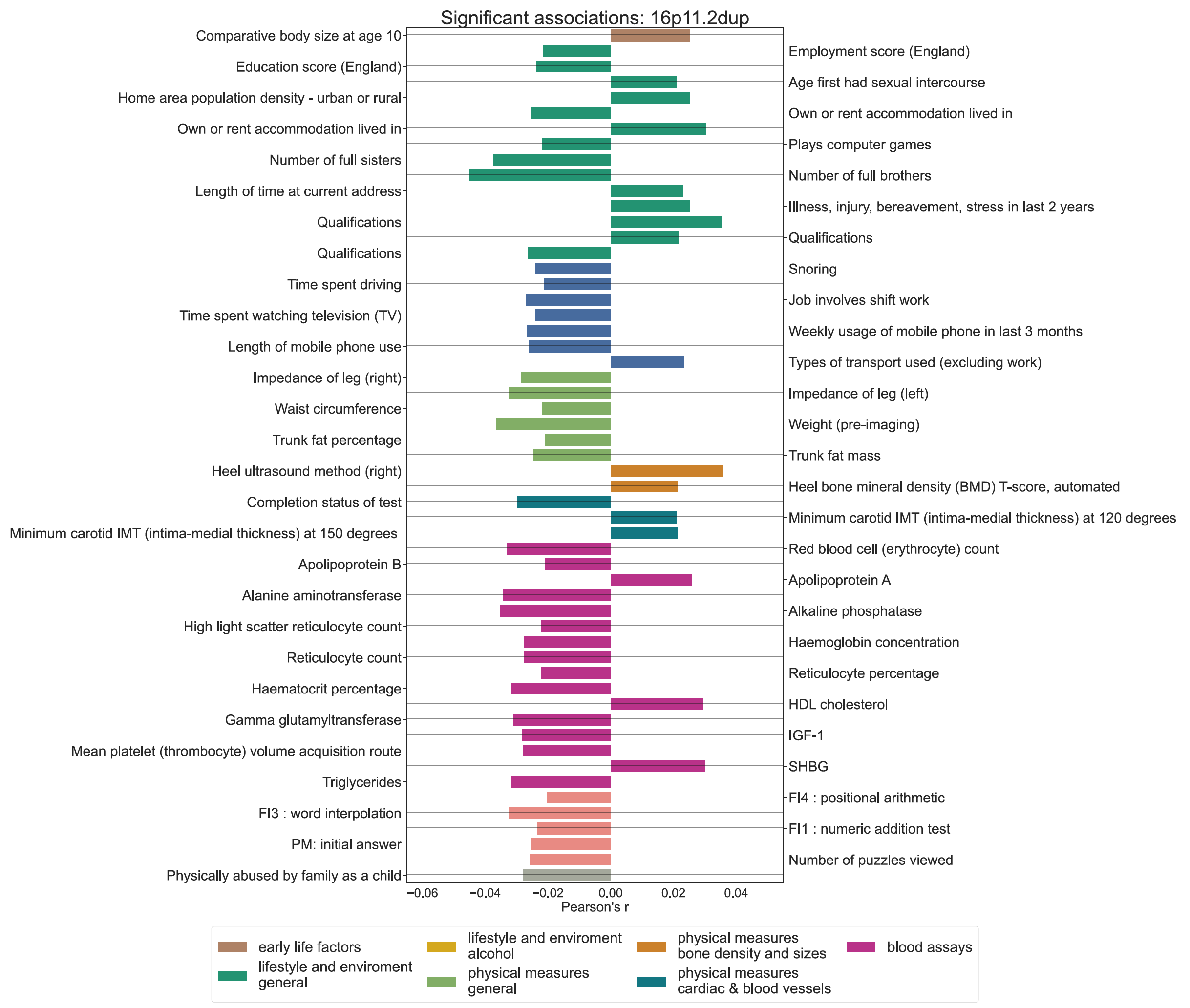

**Supplementary Figure 8**

Significant PheWAS associations for 16p11.2 proximal duplication intermediate phenotype expression.

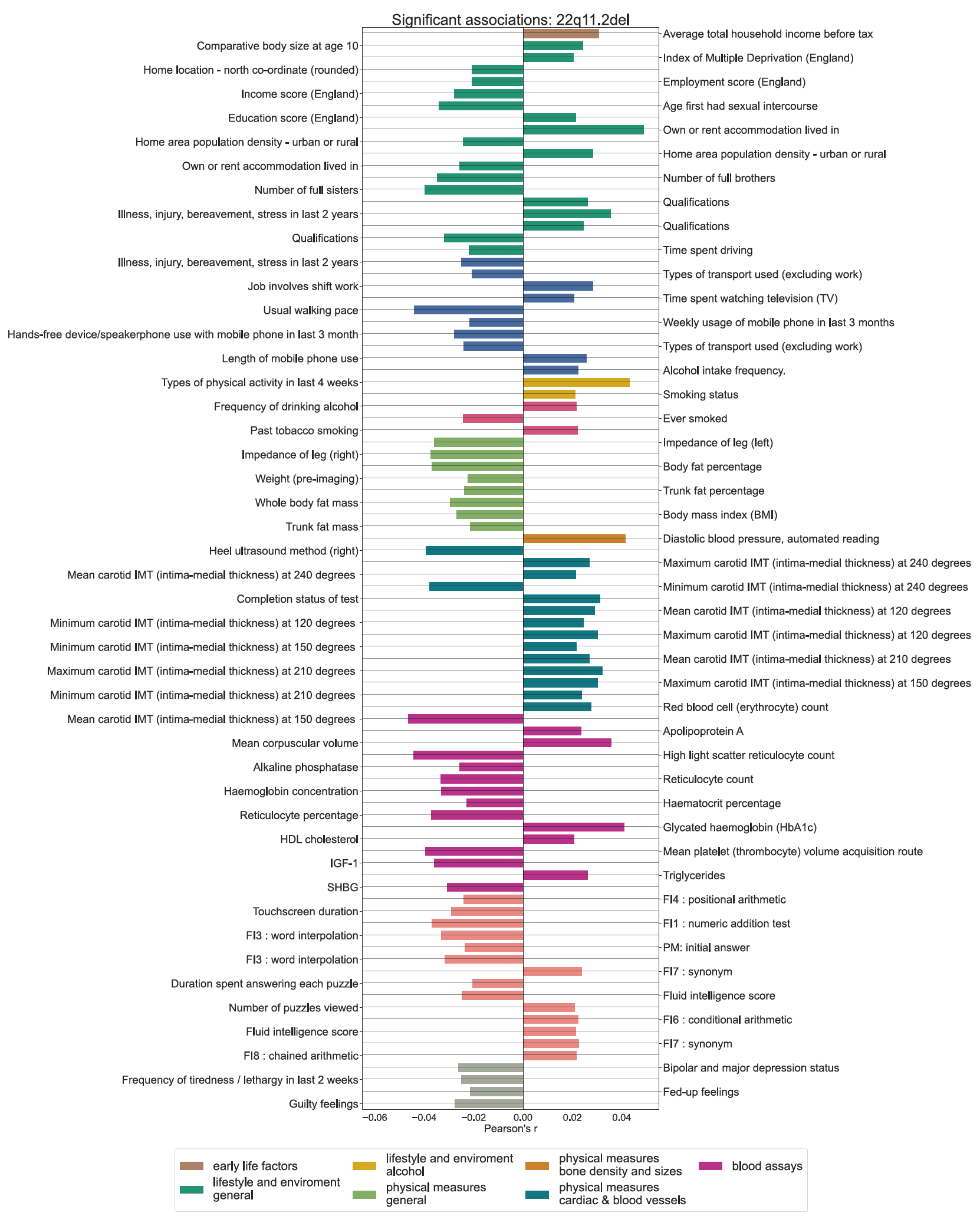

**Supplementary Figure 9**

Significant PheWAS associations for 22q11.2 deletion intermediate phenotype expression.

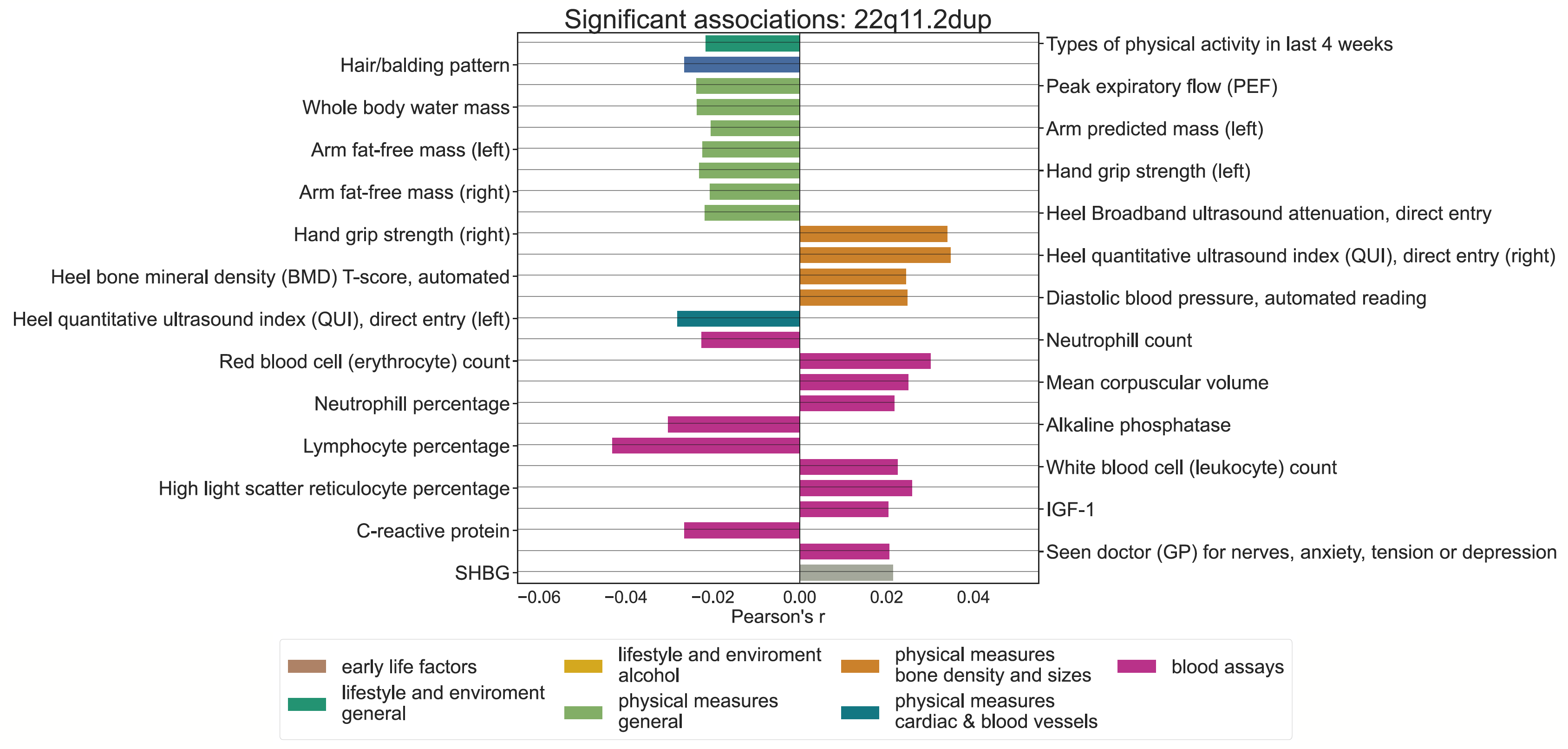

**Supplementary Figure 10**

Significant PheWAS associations for 22q11.2 duplication intermediate phenotype expression.
